# Elucidation of Structural Mechanism of ATP Inhibition at the AAA1 Subunit of Cytoplasmic Dynein 1 Using a Chemical “Toolkit”

**DOI:** 10.1101/2021.06.11.448084

**Authors:** Sayi’Mone Tati, Laleh Alisaraie

## Abstract

Dynein is a cytoskeletal motor protein that carries organelles via retrograde transport in eukaryotic cells. The motor protein belongs to the ATPase family of proteins associated with diverse cellular activities and plays a critical role in transporting cargoes to the minus end of the microtubules. The motor domain of dynein possesses a hexameric head, where ATP hydrolysis occurs. The AAA1 binding site is the leading ATP hydrolytic site, followed by the AAA3 subsite. Small-molecule ATP competitive inhibitors of dynein are thought to target the AAA1 site. The presented work elucidates the structure-activity relationship of dynapyrazole A and B, ciliobrevin A and D in their various protonated states and their 46 analogs for their binding properties in the nucleotide-binding site of the AAA1 subunit and their effects on the functionally essential subsites of the motor domain of cytoplasmic dynein 1, as there is currently no similar experimental structural data available. Ciliobrevin and its analogs bind to the ATP motifs of the AAA1, namely the Walker-A or P-loop, the Walker-B, and the sensor I and II. Ciliobrevin A shows a better binding affinity to the AAA1 binding site of dynein 1 than its D analog. Although the double bond in ciliobrevin A and D was expected to decrease the ligand potency, they show a better affinity to the AAA1 binding site than dynapyrazole A and B, lacking the bond. Protonation of the nitrogen in ciliobrevin A, D, dynapyrazole A, and B at the N9 site of ciliobrevin, and the N7 of the latter increased their binding affinity. Exploring ciliobrevin A geometrical configuration suggests the *E* isomer has a superior binding profile over the *Z* due to binding at the critical ATP motifs. Utilizing the refined structure of the motor domain obtained through protein conformational search in this study exhibits that Arg1852 of the yeast cytoplasmic dynein could involve in the “glutamate switch” mechanism in cytoplasmic dynein 1 in lieu of the conserved Asn in AAA+ protein family, as the guanidine moiety of the Arg engages in an H-bond with the carboxylate moiety of Glu1849.

## 1. Introduction

Motor proteins, dynein, kinesin, and myosin, in eukaryotic cells are responsible for transporting cargoes within cells [1]. Dynein and kinesin perform their function in conjunction with the cytoskeletal protein microtubule (MT) [1]. MTs are comprised of 11-16 protofilaments biopolymers [2] consisting of αβ-heterodimer proteins [1, 3]. Nine subfamilies compose the dynein family: namely, seven axonemal and two cytoplasmic, dynein 1 and dynein 2 [4]. Cytoplasmic dynein 1 drives retrograde axonal transport [5] and also plays a role in the mitosis process of cell division [1]. Cytoplasmic dynein 2 guarantees transportation of cargoes through MTs in flagella, motile and primary cilia [6, 7]. This isoform is also referred to as Intraflagellar Transport (IFT) dynein [6, 7]. IFT is critical to the Hedgehog pathway (Hh pathway), which refers to an essential mediator during the development of the embryo and oncogenesis [8]. It facilitates anterograde and retrograde trafficking of transcription factors such as Gli1 and Gli2 during the Hh-pathway [9, 10]. Impairment of dynein 2 could disturb the Hh pathway since it is involved in IFT [6, 7]. Inhibitors of dynein such as ciliobrevin analogs cause the inhibition of the Hh pathway [10]. Malfunction of dynein can promote cancerous cell proliferation [7], as dynein 2 is involved in the Hh-pathway and oncogenesis process [6].

Defects in the heavy chain of dynein are associated with neurodegenerative diseases (NDDs) [11], characterized by the degradation of neurons. NDDs refers to an array of neurological disorders, including Parkinson’s disease (PD), Huntington’s disease (HD), Alzheimer’s disease (AD), and motor neuron diseases [5]. Three common features observed in the NDDs are the presence of protein aggregates, the involvement of non-autonomous factors, and the dysfunction in axonal transport [5]. PD is characterized by the death of dopaminergic cell groups producing dopamine in the substantia nigra, which results in symptoms such as resting tremors, bradykinesia, and rigidity of limbs [5]. HD is a condition associated with disturbance in muscle coordination and cognitive impairment caused by a polyglutamate fragment on the huntingtin protein resulting from the repetition of the CAG codon in exon 1 of the gene responsible for the mentioned protein [5, 11]. Both PD and HD affect basal ganglia in the brain [5]. The occurrence of axonal dystrophy in the brain of patients with PD indicates abnormalities in axonal transport. Dysfunction of the axonal transport observed in animal, and cellular models represent indirect evidence of dynein involvement in PD and HD pathologies. Dysfunction of dynein causes the Golgi apparatus to fragmentize, a phenomenon observed in the brain of patients with PD, cellular and animal models of PD and HD [5]. AD, affecting 25 million individuals globally, is characterized by progressive deterioration of memory that results in this pathology and the loss of cognitive abilities, poor judgment, and speech impairment [11]. AD is marked by the presence of clusters of misfolded proteins, amyloid plaques consisting primarily of amyloid β peptides (Aβ) in the brain of patients with AD [5]. Indirect evidence obtained through the knock-down of dynein, causing an increase of Aβ peptides, suggested the involvement of dynein in AD; however, it accounts for further experiments to exhibit a direct correlation between dynein activity and AD [5]. Eyre *et al*. revealed that dynein plays an essential role in the transportation of NS5A, a hepatitis C viral protein, inside cells. Dynein ensures the efficient replication of the virus as well as the assembly of virions [6]. Dynein was discovered before kinesin [4]; however, the former has been more challenging than the latter to solve its three-dimensional structure and to understand the exact mechanism of action of its multi-domain construction. Indeed, the complexities result from its massive size, its two heavy chains of each 530 kDa[7]. Despite the complexity of dynein structure, characterization of some of its substructures or domains utilizing X-ray crystallography, electron microscopy, and mutagenesis studies have provided insights into understanding its function and role in cell [7]. Dynein is a homodimer protein [1], composed of two heavy chains (HCs), each 530 kDa; two light intermediate chains (LICs), each 74 kDa; four intermediate chains (IC) with weight varying between 53 kDa and 59 kDa each, and six light chains (LCs), each 10-14 kDa [11]. The heavy chain of dynein consists of the tail, the linker, the hexameric head, the buttress, the stalk domains, and the microtubule-binding domain (MTBD) [12, 13]. The N-terminus of dynein, representing approximately a third of the 530 kDa heavy chain, constitutes the tail and the linker [1, 14]. **(Figure 1)**

**Figure 1:**
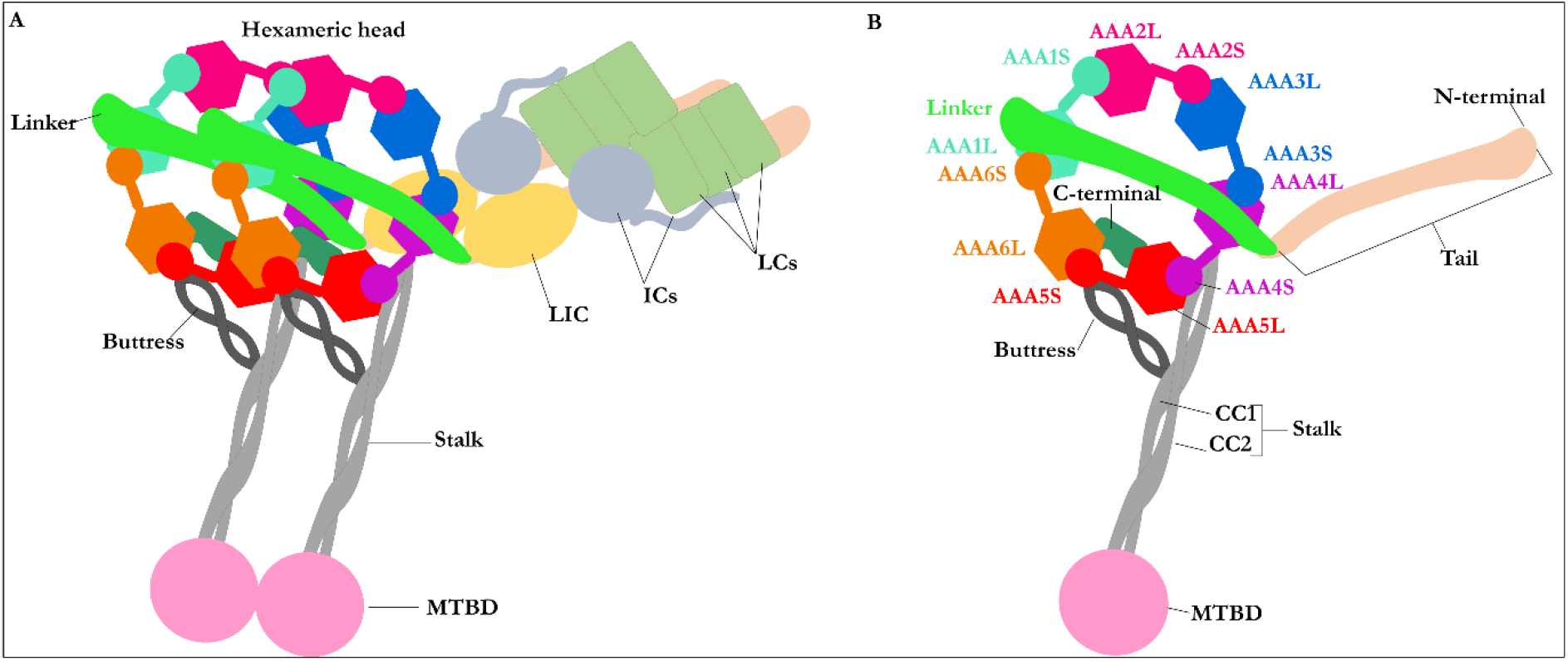
The multi-domain structure of cytoplasmic dynein. **A)** Schematic representation of the homodimer cytoplasmic dynein. For simplicity, only one monomer is labeled. The homodimer represents two heavy chains, two LICs, four ICs, and six LCs. Each set of the two heavy chains consists of a tail, a linker, a hexameric head, a buttress, a stalk, and an MTBD. The figures (e.g., the LCs, ICs, and LICs) are schematic. They do not represent their actual shape, **B)** Schematic representation of the heavy chain of cytoplasmic dynein in its post-Powerstroke conformation, with the linker straight and positioned on AAA4 near the stalk. The hexameric head represents the six AAA+ subunits with small (AAA+S) and large (AAA+L) subunits.

The tail of the heavy chain of dynein represents the site for cargo binding, where the dimerization of both monomers occurs [1]. The tail also binds to the LICs and ICs [1, 5]. The linker domain follows the tail and is thought to be involved in a force-generating process as its position changes upon binding of ATP, resulting in the motility of dynein [1]. The stalk of ∼10-15 nm in length is attached to the MTBD at the C-terminus [15]. It is linked to and supported by buttress [1]. **(Figure 1 A)**

The head or motor domain of dynein is comprised of six AAA+ subunits, four of which (AAA1 to AAA4) possess a nucleotide-binding site at the interface between one subdomain and the subsequent subdomain; the AAA1 nucleotide-binding site is enclosed between the AAA1 and AAA2 subdomains [1, 12]. Three of the four binding sites, AAA1, AAA3, and AAA4, present the ability to hydrolyze ATP [1, 12]. Each of the six AAA+ subdomains encompasses a small and a large subunit linked by a flexible unfolded segment [1]. **(Figure 1 B & Figure 2 B)**

**Figure 2:**
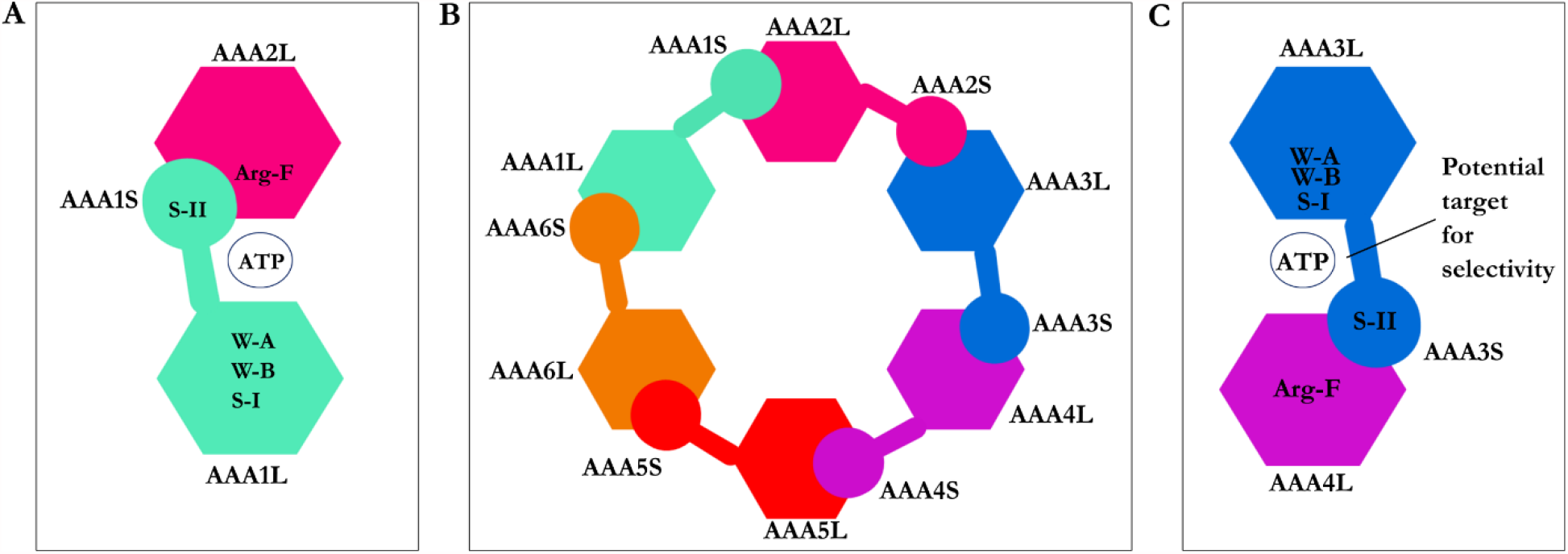
Composition of the AAA+ subdomains. **A)** The AAA1 nucleotide-binding site composed of the small (AAA1S) and large (AAA1L) subunits of AAA1 and the prominent (AAA2L) subunit of AAA2. ATP motifs are represented: walker-A (W-A), walker-B (W-B), sensor I (the S-I), sensor II (the S-II) in AAA1, and Arginine finger (Arg-F) in AAA2, **B)** Hexameric head of cytoplasmic dynein, **C)** The AAA3 nucleotide-binding site composed of its small (AAA3S) and large (AAA3L) subunits and the large subunit of AAA4. ATP motifs are represented as the walker-A (W-A), the walker-B (W-B), the sensor I (the S-I), the sensor II (the S-II) in AAA3, and the Arginine finger (Arg-F) in AAA4.

The AAA1 is the primary site of ATP hydrolysis in cytoplasmic dynein [1] since hydrolysis of ATP at this site is critical for dynein motility [16] and conserved in dynein family [1]. The AAA3 is the second major site of ATP hydrolysis [1], as mutation of K2675T in the D. *discoideum* species reduced ATPase activity of dynein by approximately 20-fold [16]. The nucleotide-binding sites of cytoplasmic dynein, similar to other AAA+ family members, display the ATP motifs: the walker-A (GXXXGK) or P-loop, the walker-B (catalytic Asp and Glu), the S-I (Asn), the S-II (Arg), an Arginine finger (Arg) and directly interacting amino acids with the nucleotide base [1, 7]. **(Figures 2-3 and Table 3)**

Considering the vital role of the heavy chain defects in causing some of the major NDDs [11] and the gigantic size of dynein (∼ 1.5 MDa), small-molecule inhibitors are suitable means to examine how the function of the dynein motor domain could be regulated or inhibited. Therefore, this structure-activity relationship (SAR) study attempted to elucidate the structural effect of ciliobrevin A, D and their analogs on their potential regulatory or inhibitory [6] mechanisms concerning the function of the motor domain in cytoplasmic dynein 1. As the size and the complexity of the various structural domains of dynein make considerable challenges to solve their atomistic *holo* or *apo* structure *in vitro, in silico* methods in the presented work were utilized to address some of the current shortcomings.

## 2. Materials and Methods

### 2.1. Structure of Dynein Motor Sub-domains

Three crystal structures of the motor domain of cytoplasmic dynein 1, including the linker, are available in the Protein Data Bank (PDB) [17]. The three crystal structures studied here are motor domains of *Dictyostelium*-motor-ADP (3VKG.pdb) [18] from *Dictyostelium discoideum*, yeast-motor-*apo* (4AKG.pdb) [19], and yeast-motor-AMPPNP (4W8F.pdb) [20], both from *Saccharomyces cerevisiae*. The hexameric head from *Dictyostelium*-motor-ADP [18] crystal structure accommodates one ADP molecule in the AAA1, AAA2, AAA3, and AAA4 subunits (3VKG.pdb) [18]. The hexameric head from the yeast-motor-AMPPNP crystal structure (4W8F.pdb) [20] possesses an AMPPNP molecule in each of the subunits. The yeast-motor-*apo* crystal structure (4AKG.pdb) [19] presents the AAA1 binding site in its unliganded state, whereas an ATP is found in the AAA2 and an ADP in the AAA3 binding site (4AKG.pdb) [19]. The three crystal structures have their linker in the post-Powerstroke conformation. The linker is straight, in the *Dictyostelium*-motor-ADP [18] crystal structure (3VKG.pdb) [18], and spans the AAA1 to AAA5 subunits. In comparison, the linker stretches from the AAA1 to the AAA4 in the yeast-motor-AMPPNP (4W8F.pdb) [20] and the yeast-motor-apo (4AKG.pdb) [19] crystal structures. The AAA1 binding sites of yeast-motor-AMPPNP [20] and *Dictyostelium*-motor-ADP [18] are in *holo* states in their crystal structures, with AMPPNP and ADP bind in each, respectively. **(Table 1)**

**Table 1:**
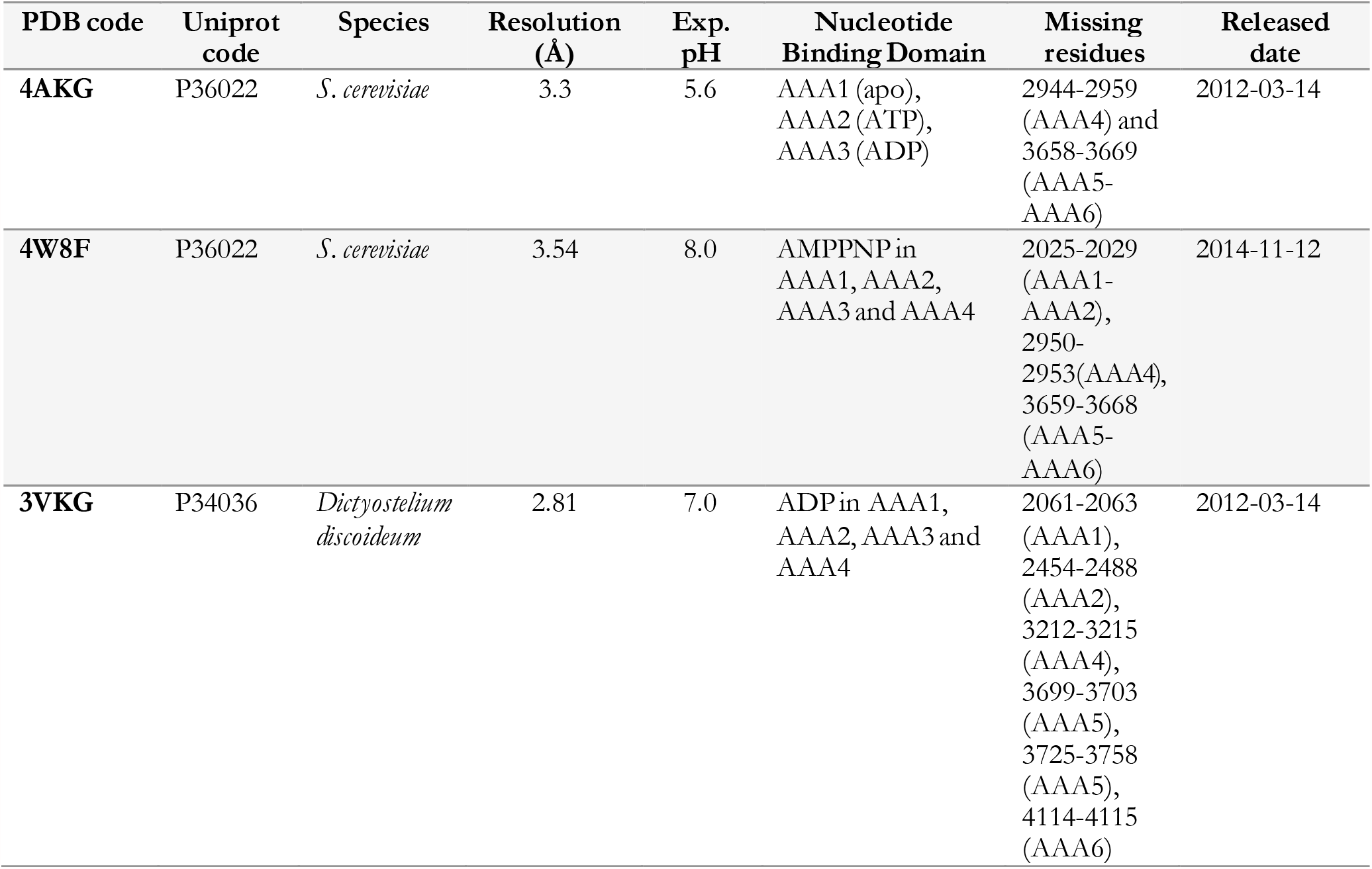
Summary of the features of the crystallographically solved structures of dynein used in this study.

### 2.2. Protein Structure Preparation

The accession codes of the three-dimensional structure of the motor domain of dynein collected from the PDB platform are 4AKG.pdb [19] from *S. cerevisiae* (Uniprot P36022 [21]), 4W8F.pdb [20] from *S. cerevisiae* (Uniprot P36022 [21]), and 3VKG.pdb [18] (Uniprot P34036 [21]) from *Dictyostelium Discoideum* (Uniprot P34036 [21]). **(Table 1)**

The amino acid sequences of the two species, *S. cerevisiae* (P36022) and *D. discoideum* (P34036), were aligned according to the ClustalW algorithm [22, 23]. There is a sequence identity of ∼25% between the cytoplasmic dynein of *S. cerevisiae* (P36022) containing 4092 residues and *D. Discoideum* (P34036) containing 4730 residues. According to the sequence alignment (Tyr1758-Val2273: *S. cerevisiae* and Tyr1936-Leu2531: *D. discoideum*), there are 35% conserved residues within the AAA1 and AAA2 subdomains of both species. **(Figure S1 & Table 2)**

**Table 2:**
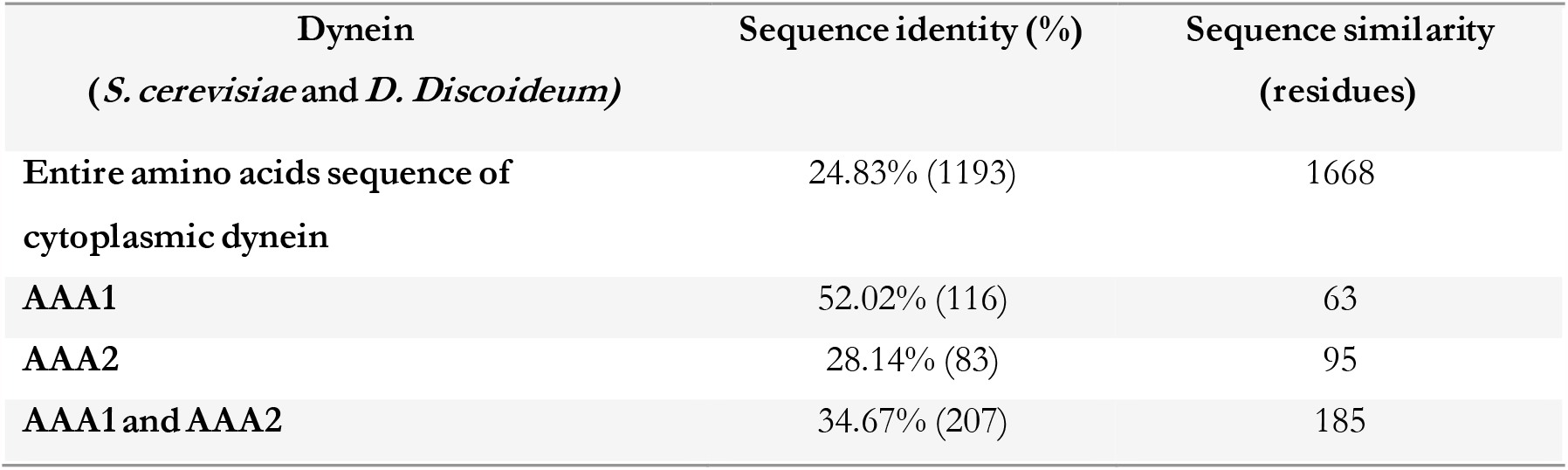
Amino acids sequence identity and similarity of the different parts of dynein, between *S. cerevisiae* and *D. discoideum*.

The yeast-motor-AMPPNP crystal structure (4W8F.pdb) [20] was subjected to E1849Q mutation to prevent ATP hydrolysis at the AAA1 nucleotide-binding site [20]. The yeast-motor-AMPPNP crystal structure was considered suitable for docking ATP competitive inhibitors in the AAA1 nucleotide-binding site and corresponded to the conformation of dynein before ATP hydrolysis [20]. The *Dictyostelium*-motor-ADP (3VKG.pdb) possesses a molecule of ADP in the AAA1 nucleotide-binding site corresponding to the configuration succeeding ATP hydrolysis [18]. In contrast, the yeast-motor-*apo* (4AKG.pdb) pertains to motor domain conformation with low-affinity nucleotides binding [19]. Thus, the yeast-motor-AMPPNP (4W8F.pdb) [20] conformation was chosen over the *Dictyostelium*-motor-ADP (3VKG.pdb) [18] or the yeast-motor-apo (4AKG.pdb) [19] for ligand docking experiment.

The missing residues (i.e., crystallographically unsolved) from the motor chain A in 4W8F.pdb [20] (i.e., Ala2025-Leu2029, Lys2950-Val2953, Lys3659-Arg3668) were modeled and completed based on the primary structure of the cytoplasmic dynein heavy chain of *S. cerevisiae* (P36022). **(Table 1)** Fourteen (14) residues from AAA1 and AAA2 subunits are located in the nucleotide-binding site. They consisted of the W-A or the P-loop region GPAGTGKT [4, 7, 18] (Gly1796-Thr1803 in *S. cerevisiae* and Gly1974-Thr1980 in *D. discoideum*), the W-B region [4, 7, 18] (Asp1848 and Glu1849 in *S. cerevisiae* compared to Asp2026 and Glu2027 in *D. discoideum)*, the S-I [4, 7, 18] (Asn1899 in *S. cerevisiae* and Asn2078 in *D. discoideum*), the S-II (Arg1971 in *S. cerevisiae* and Arg2150 in *D. discoideum)*, the Arg finger (Arg2209 in *S. cerevisiae* and Arg2410 in *D. discoideum)* [4, 7, 18], and the N-loop [4, 7, 18] (Leu1769 and Ile1770 in *S. cerevisiae* compared to Leu1947 and Val1948 in *D. discoideum)*. **(Table 3)**

**Table 3:** The ATP motifs in *S. cerevisiae* and *D. discoideum*.

| ATP Motifs | *S. cerevisiae* | *D. discoideum* |
| --- | --- | --- |
| **Walker-A** | Gly1796-Thr1803 | Gly1974-Thr1980 |
| **Walker-B** | Asp1848-Glu1849 | Asp2026-Glu2027 |
| **Sensor I** | Asn1899 | Asn2078 |
| **Sensor II** | Arg1971 | Arg2150 |
| **Arg finger** | Arg2209 | Arg2410 |
| **N-loop** | Leu1769-Ile1770 | Leu1947-Val1948 |

The retrieved X-ray crystal structure (4W8F.pdb) [20] was truncated to keep the required domains potentially affecting the nucleotide-binding sites to reduce the necessary CPU time for the motor sub-domains conformational search. That reduced the number of atoms for calculating bonding and non-bonding interactions among ligand and the protein atoms. The resulting truncated structure included, dynein hexameric head (AAA1-AAA4: Tyr1758-Val2984 and AAA5-AAA6: Leu3370-Asn3970), the linker subunit (within the tail: Gly1363-Gln1757), and a part of the stalk (Ile2993-Ser3125) interacting with the hexameric head. GROMACS [24] package v. 2016.5, with Gromos 96 force field 54A7 [25], was utilized for generating topology of protein atoms and energy minimization *in vacuo* to optimize bond lengths, angles, and orientation of the residues in the protein structure before docking any ligands.

### 2.3. Ligands 3D Structure Preparation

The AMPPNP’s atomic coordination at the AAA1 site (4W8F.pdb) [20] was used as the reference. The binding site region was specified at a 15.0 Å radius spherical region around the reference structure as the center, covering an extra 2.0 Å broader region than that occupied by the AMPPNP interacting - amino acids in the binding site of the AAA1 domain of cytoplasmic dynein 1. **(Figure 3)**

**Figure 3:**
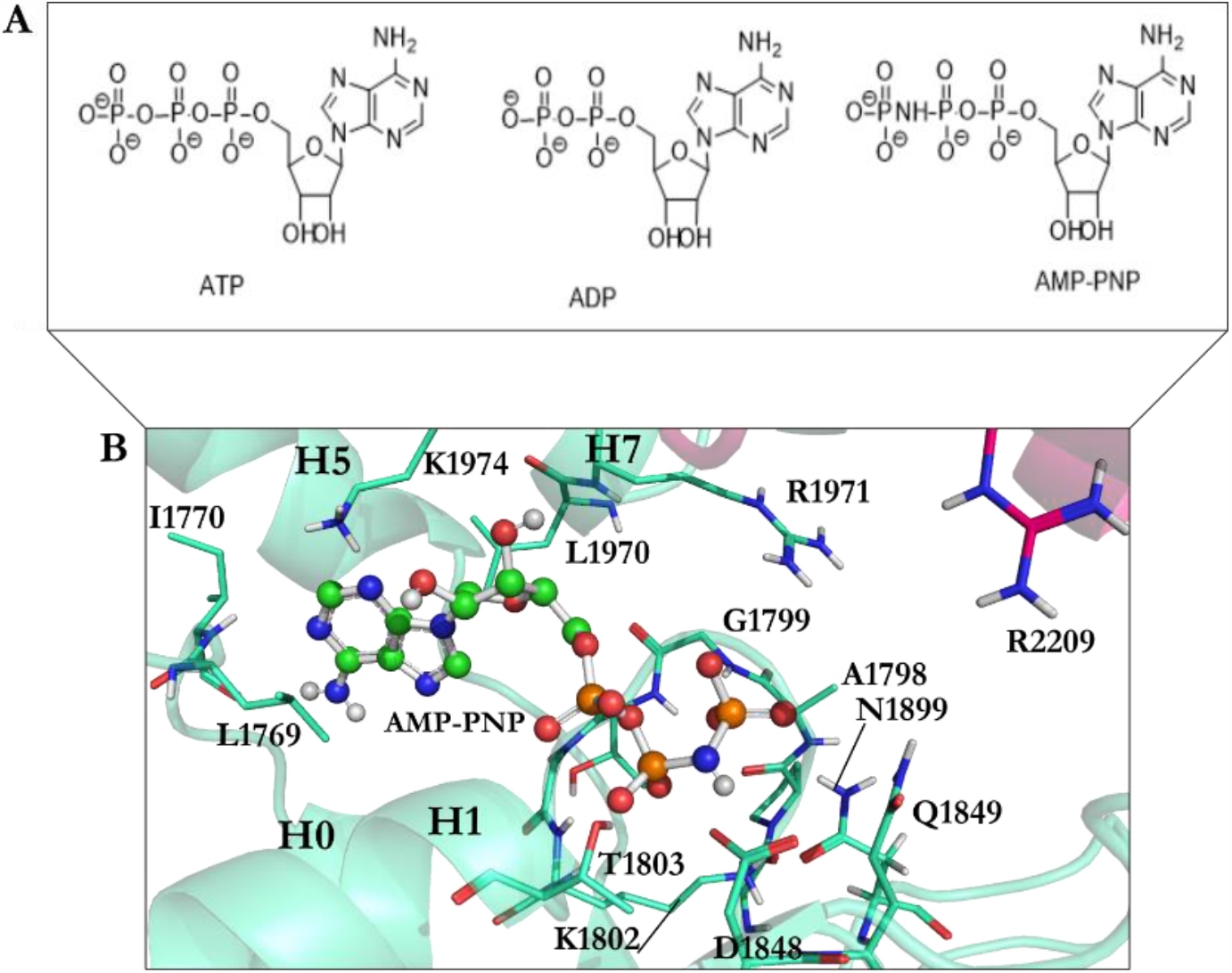
**A)** The chemical structure of ATP, ADP, and AMPPNP, **B)** AMPPNP is in the ball-and-stick representation (green) interacts with amino acids in the AAA1 binding site of cytoplasmic dynein 1. Color code: AAA1 in cyan blue and AAA2 in sharp pink.

A library of 63 ligands (i.e., a chemical tool kit in this study) was created using SYBYL-X 2.1.1 (Cetara Corporation©). Three-dimensional structures of the ligands were built up individually and minimized stepwise using the steepest descent algorithm according to the Tripos force field, with 0.0001 kJ/mol energy gradient and 10,000,000 iterations. The library contained previously synthesized and *in vitro* studied 46 analogs of ciliobrevin [10], dynapyrazole A and B [26], and the protonated forms of the lead compounds ciliobrevin A and D, dynapyrazole A and B modeled *in silico*. It also included the nucleotides ATP, ADP, and AMPPNP, a non-hydrolyzable analog of ATP. **(Figures 3-5)**

ATP and ADP are the endogenous substrates of dynein 1 [1]. Since the crystal structure of yeast-motor-AMPPNP did not possess ATP or ADP in any of the four nucleotide-binding sites (AAA1-AAA4), the endogenous substrates were docked into the AAA1 binding site to study their binding mode and quantify the magnitude of their binding affinity versus that of each ligand in the library. The deprotonated form of ATP, ADP, and AMPPNP, was based on the ATP pKa values [27]. The pH was 8.0 during the crystallization of the yeast-motor-AMPPNP(4W8F.pdb) [20], and the pKa of γ-phosphate is approximately 6.49 [27]. In comparison, the pKa of the α- and β-phosphates of ATP are estimated at ∼ 1.6 [27]. Therefore, ATP, ADP, and AMPPNP molecules were also built and assessed in their fully deprotonated state and subjected to energy minimization. The protonated ciliobrevin A and D structures, dynapyrazole A and B were built up and energetically minimized. The pKa of the inhibitors have not yet been experimentally defined. A study on the different components of the ligands’ chemical structures helped to study the effect of the most probable protonation states on their binding affinity. The pKa of arylamine groups, existing in the structures of the ligands, varies between 9-10 [28], meaning that at the physiological pH, an arylamine (i.e., consisting of the N9 atom of dynapyrazole A and B, ciliobrevin A and D, and their analogs) could be protonated. Noteworthy that the lone-pair electrons of the N7, N9, and N11 in dynapyrazole A and B, could be involved in delocalized electronic systems of A, B, and C fragments, reducing the availability of the lone-pair electrons for protonation. Furthermore, the pKa of quinazoline-4(3*H*)-one moiety of dynapyrazole (i.e., ring A and B) is expected to be more acidic than the estimated 3.51 of quinazoline [29], due to the electron withdrawal effect of the oxygen. Thus, the moiety is more likely to be deprotonated at the physiological pH. **(Figure 4)**

**Figure 4:**
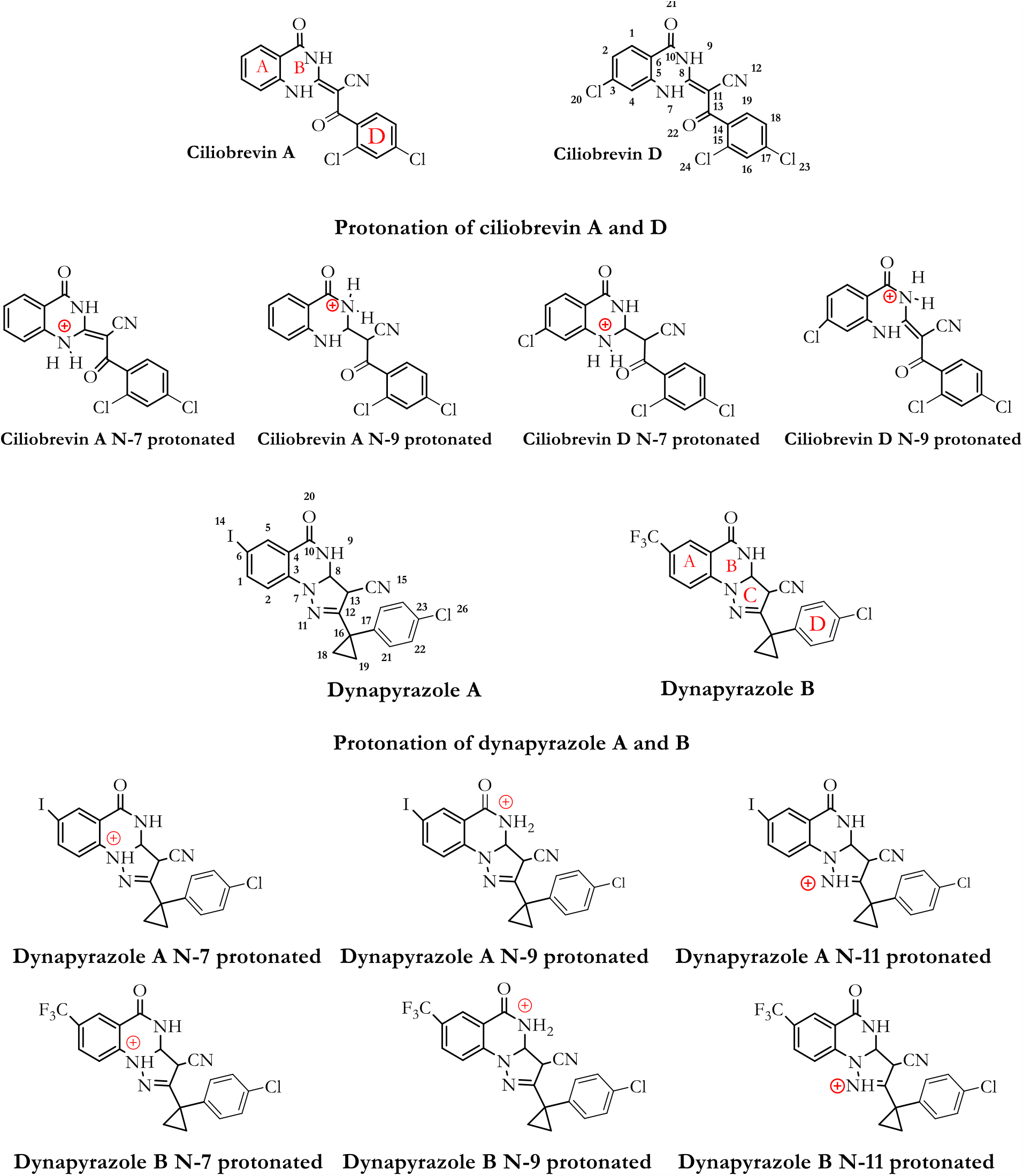
Chemical Structure of ciliobrevin A and D, dynapyrazole A and B, as well as their protonated structures in the ligand library. Ciliobrevin A and D are respectively analog 1 and 2. Chemical structures and atom numbering were obtained utilizing Chemdraw software.

FlexX [30, 31] docking software, embedded in the LeadIT software package (v.2.1.8), was utilized for ligand-protein binding mode predictions, energy estimation, and ranking the solutions. It predicts the protein-ligand interactions based on the incremental construction algorithm [32]. There are three fundamental stages to the FlexX docking algorithm: selecting a base fragment, placing the base fragments into the active site, and incrementally constructing the complex, and calculating the interaction energies according to the Böhm scoring function for ranking the docking solutions [33, 34].

## 3. Results and Discussion

### 3.1. Dynapyrazole, Ciliobrevin and their Analogs

*In vitro* and *in vivo* studies of ciliobrevin A and D, the two ATP-competitive ligands, have shown that they non-selectively bind to the ATP binding sites of the hexameric head of both cytoplasmic dynein 1 and dynein 2 [7, 10]. Dynapyrazole A and B resulted from a chemical structure modification to produce ciliobrevin analogs with higher potency [26], for overcoming geometric isomerization complexity caused by the C8-C11 double bond in ciliobrevin. **(Figure 4)**

Unlike the ciliobrevin analogs, which abrogate both MT-stimulated and basal ATPase activity, dynapyrazole analogs inhibit MT-stimulated ATPase activity with high potency without affecting basal ATPase activity [26]. This feature resembles She1, a microtubule-associated protein (MAP) that effectively reduces MT-stimulated ATPase activity without significantly decreasing its basal activity [35]. Experiments have shown that ciliobrevin A and D, which bind to AAA1, might bind to the AAA3 site [10]. In contrast, analogs of dynapyrazole, especially compound 20, abolished basal dynein activity by binding to the AAA3 and AAA4 [36]. Forty-six (46) analogs of ciliobrevin A and D were proposed to have potentially higher selectivity and potency than ciliobrevin A against dynein 2 [7]. However, only the IC_50_ of four analogs (i.e., 18, 37, 43, and 47) against dynein 1 and 2 were reported [7]. Their structural binding modes were investigated as follows. **(Figure 5 & Table 4)**

**Table 4:** IC_50_ of ciliobrevin A and D, their analogs 18, 37, 43, and 47, dynapyrazole A and B for dynein 1 and dynein 2. *\* The IC*_*50*_ *of dynapyrazole B against dynein 1 is not available*.

| Compounds | $IC_{50}$ (μM) | $IC_{50}$ (μM) |
| --- | --- | --- |
|  | Dynein 1 | Dynein 2 |
| Ciliobrevin A | 52.0 | 55.0 |
| Ciliobrevin D | 15.0 | 15.5 |
| Dynapyrazole A | 2.3 | 2.6 |
| Dynapyrazole B* | – | 2.9 |
| Analog 18 | 130.0 | 21.0 |
| Analog 37 | 280.0 | 11.0 |
| Analog 43 | 158.0 | 16.0 |
| Analog 47 | 130.0 | 11.0 |

**Figure 5:**
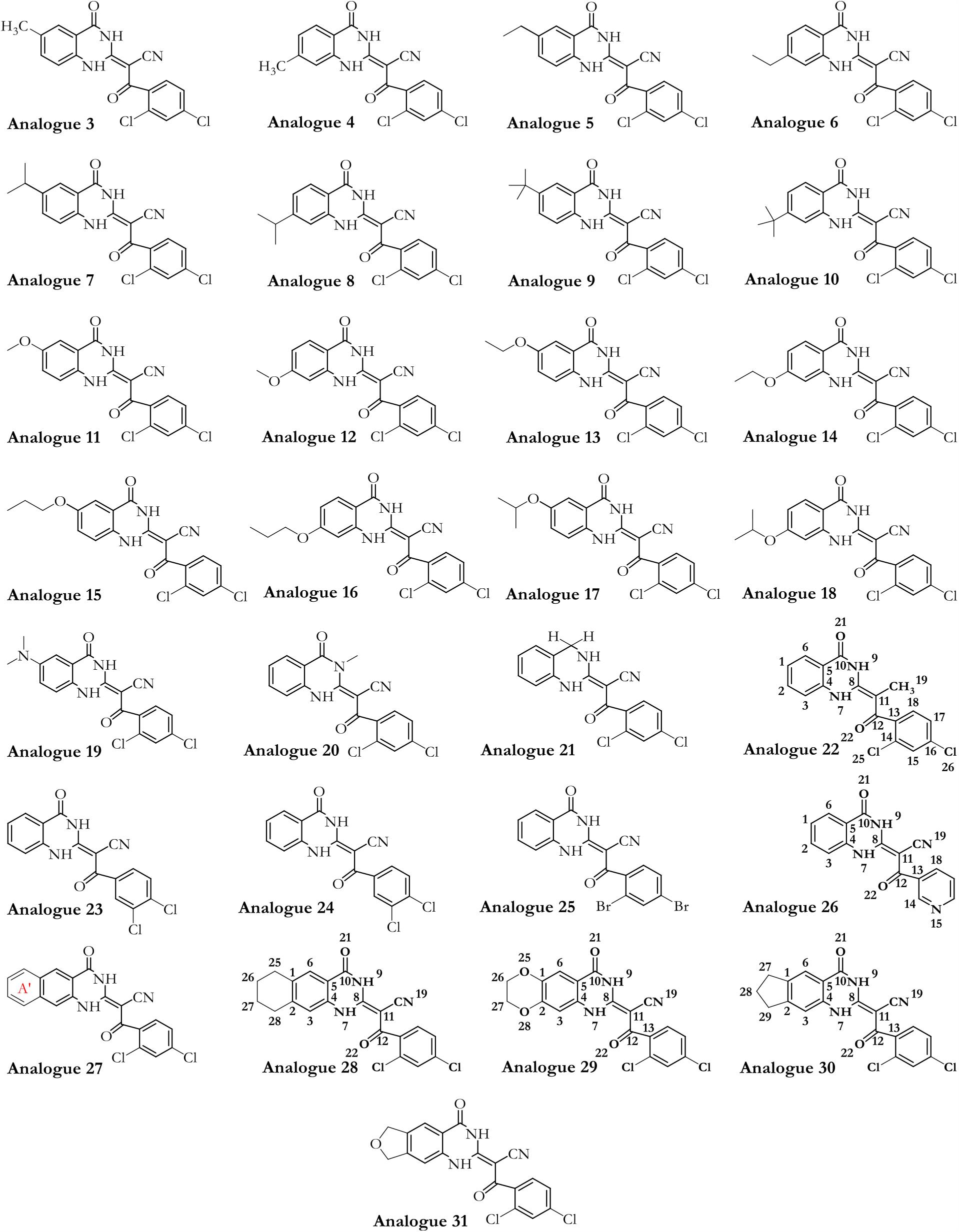
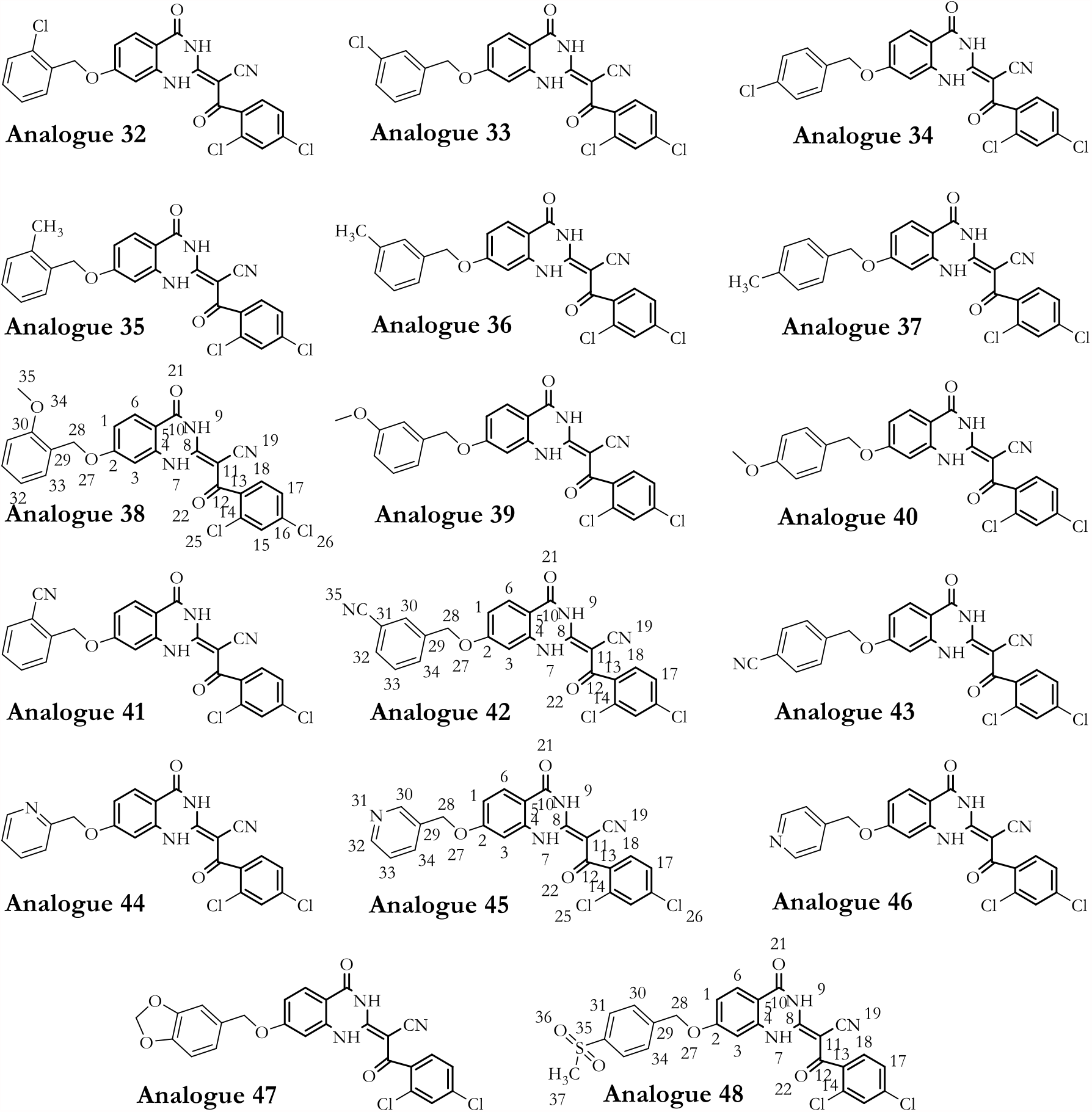
Forty-six (46) analogs (from analogs 3 to 48) of ciliobrevin in the ligand library. Chemical structures and atom numbering were obtained utilizing Chemdraw software.

### 3.2. Dynapyrazole, Ciliobrevin, and their Analogs Binding

Docking of the AMPPNP, obtained from the crystal structure, into the binding site of the yeast-motor-AMPPNP [20] of dynein resulted in a conformation with the lowest RMSD (1.67 Å) and binding energy (−22.06 kJ/mol). The ligand interacted with: the N-loop (Pro1766-Leu1774) via residues Leu1769 and Ile1770, the W-A region (Gly1799-Thr1803), the β6 strand including S-I motif (Ala1893-Asn1899) via Asn1899, Ile1929 from H5 (Ser1926-Ile1936), and Leu1970, Lys1974 from the H7 (Leu1970-Pro1982). **(Figure 6 C-D, Figure S1, and Table 5)**

**Table 5:** Binding properties of the nucleotides’ conformation obtained from the *in silico* experiments.

| Ligand | RMSD vs.<br>X-ray structure | Binding<br>energy<br>(kJ/mol) | Residues interacting with the compound |
| --- | --- | --- | --- |
| AMPPNP | 1.67 | -22.06 | Leu1769, Ile1770, Gly1799, Gly1801,<br>Lys1802, Thr1803, Glu1804, Asn1899,<br>Ile1929, Leu1970, Lys1974 |
| AMPPNP minimized<br>structure | 4.75 | -40.18 | Glu1767, Gly1799, Gly1801, Lys1802,<br>Thr1803, Glu1804, Gln1849, Asn1899,<br>Lys1974 |
| Minimized ATP | — | -42.33 | Ala1798, Gly1799, Thr1800, Gln1849,<br>Asn1851, Arg1852, Asn1899, Arg1971 |
| Minimized ADP | — | -31.89 | Ala1798, Gly1799, Thr1800, Asp1848,<br>Gln1849, Arg1852, Asn1899, Arg1971 |

**Figure 6:**
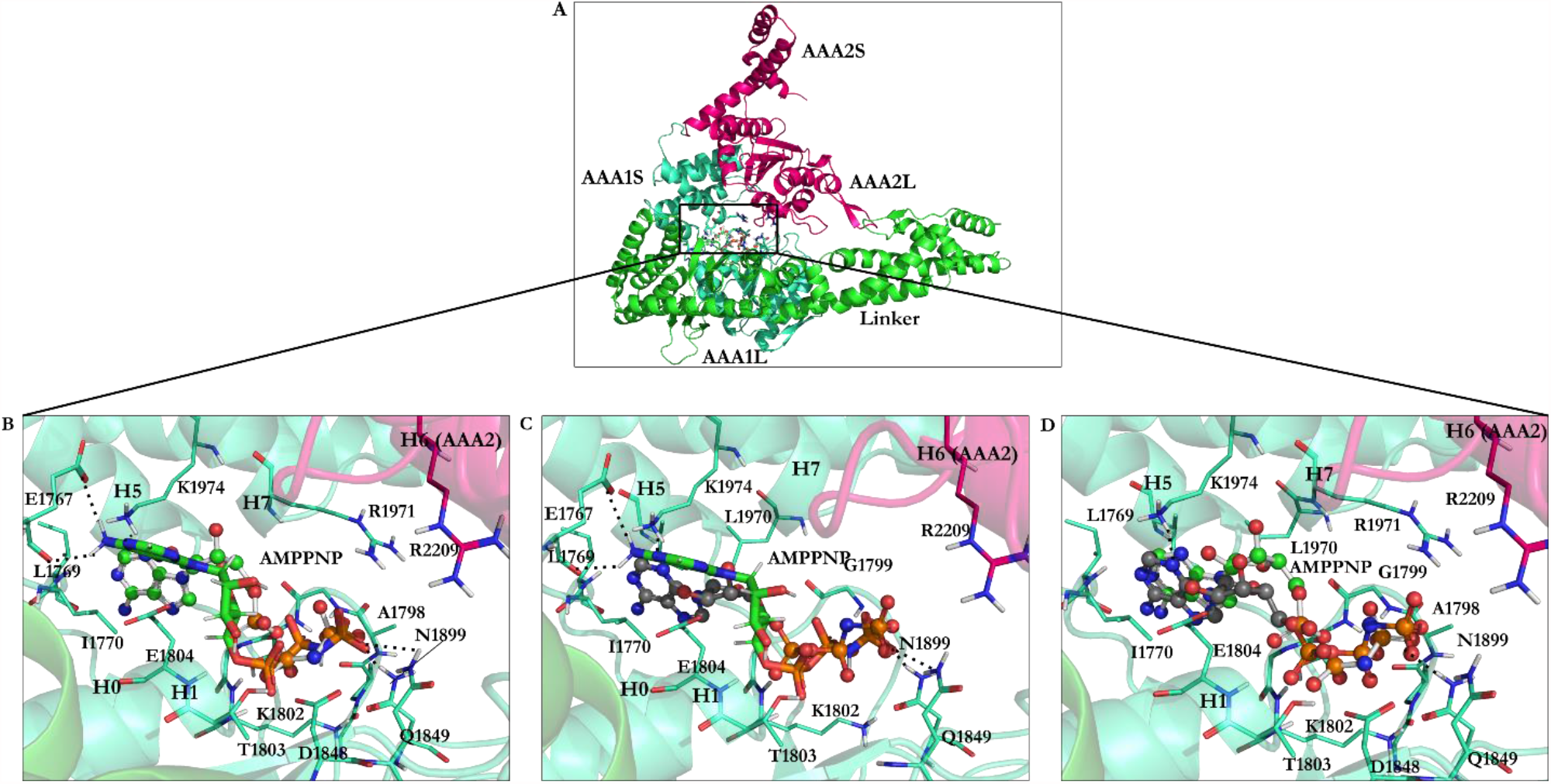
AMPPNP in the AAA1 binding site of dynein, **A)** The linker, the AAA1 and AAA2 subunits of dynein and illustration of their inter-subunit binding site, **B)** Docking solution of the minimized AMPPNP in the AAA1 binding site of the minimized conformation from yeast-motor-AMPPNP crystal structure (4W8F). The docking solution is in stick representation, the crystal structure of AMPPNP in ball and stick, **C)** The energy minimized AMPPNP in the AAA1 binding site of the energy minimized structure of yeast-motor-AMPPNP (4W8F) superimposed with the docking solution of the reference ligand, AMPPNP, from the crystal structure, **D)** Crystal structure of AMPPNP docked in the AAA1 binding site of the crystal structure of yeast-motor-AMPPNP (4W8F). The calculated conformations (black-grey) ball and stick representations and the crystal structure of AMPPNP (reference, green) ball and stick. Non-essential hydrogen atoms are not shown for simplicity.

Superposition of the domain crystal structures shows the W-A region (Gly1796-Thr1803) in the yeast-motor-apo (4AKG.pdb) [19] is ∼ 7.0 Å distant from the W-A in the yeast-motor-AMPPNP (4W8F.pdb) [20]. The W-A region (Gly1974-Thr1980 in *D. discoideum* and Gly1796-Thr1803 in *S. cerevisiae*) shifts by ∼ 1.4 Å (in *Dictyostelium*-motor-ADP) compared to that in yeast-motor-AMPPNP (4W8F.pdb) [20]. The H5 (Ser1926-Ile1936) of the AAA1 in yeast-motor-apo (4AKG.pdb) [19] crystal structure shifts by ∼ 3.8 Å from the position of the equivalent helix in the yeast-motor-AMPPNP (4W8F.pdb) [20]. The H5 helix (Arg2105-Tyr2114 in *D. discoideum* and Ser1926-Ile1936 in *S. cerevisiae*) in the *Dictyostelium*-motor-ADP (3VKG.pdb) [18] and in the yeast-motor-AMPPNP (4W8F) [20] are ∼ 1.3 Å apart, similar to the H7 (Leu1970-Pro1982) of the AAA1 in the yeast-motor-AMPPNP (4W8F.pdb) [20] and yeast-motor-*apo* (4AKG.pdb) [19] at a ∼ 3.9 Å distance. There is a ∼ 2.0 Å distance between the H7 (Gly2148-Lys2165 in D. *discoideum* and Leu1970-Pro1982 in *S. cerevisiae*) of the *Dictyostelium*-motor-ADP (3VKG.pdb) [18] and of yeast-motor-AMPPNP (4W8F.pdb) [20]. The displacements of the domain segments (i.e., yeast-motor-AMPPNP [20], yeast-motor-*apo* [19], and *Dictyostelium*-motor-ADP [18]) imply that AMPPNP binding has caused an “induced fit” driven conformational change in the binding site. **(Figures S2-S3)**

The docked AMPPNP conformation obtained from its energy minimization has a binding energy of -40.18 kJ/mol with 4.75 Å RMSD due to the conformational search and energy minimization of the ligand and the consequent optimization of bonds length and angles according to the implemented force field parameters. Similar to the reference ligand, the conformation of the docked, minimized (i.e., the optimized) structure of AMPPNP interacts with ATP motifs [4, 7]. However, the orientation of the aromatic nucleotide fragment of the energy minimized AMPPNP allowed the system to engage with positively charged Lys1974 of the H7 (Leu1970-Pro1982) via polar-ionic interactions, which is not possible for the ligand with the conformation seen in the crystal structure. Unlike the latter, the amine group of the optimized conformation engages in H-bond interactions with the carboxylate group of Glu1767 (N-loop: Pro1766-Leu1774), and its γ-phosphate creates an H-bond with Gln1849 (E1849Q). **(Figure 6 B-C, Table 3 & Table 5)**

The energy-minimized ATP’s binding energy is lower than that of AMPPNP, which suggests ATP binds more strongly to cytoplasmic dynein 1 than AMPPNP (−42.33kJ/mol vs. -40.18kJ/mol). The ATP’s binding mode obtained after the conformational search displayed its interaction with Ala1798, Gly1799, and Thr1800 from the W-A region (Gly1796-Thr1803), Gln1849 form the W-B motif in β3 (Ala1843-Asp1848), Asn1851 and Arg1852, between β3 (Ala1843-Asp1848) and H3 (Glu1854-Val1874), with the S-I (Asn1899 in β6: Ala1893-Asn1899) and Arg1971 from H7 (Leu1970-Pro1982). **(Figure 7 B, Table 3 & Table 5)**

**Figure 7:**
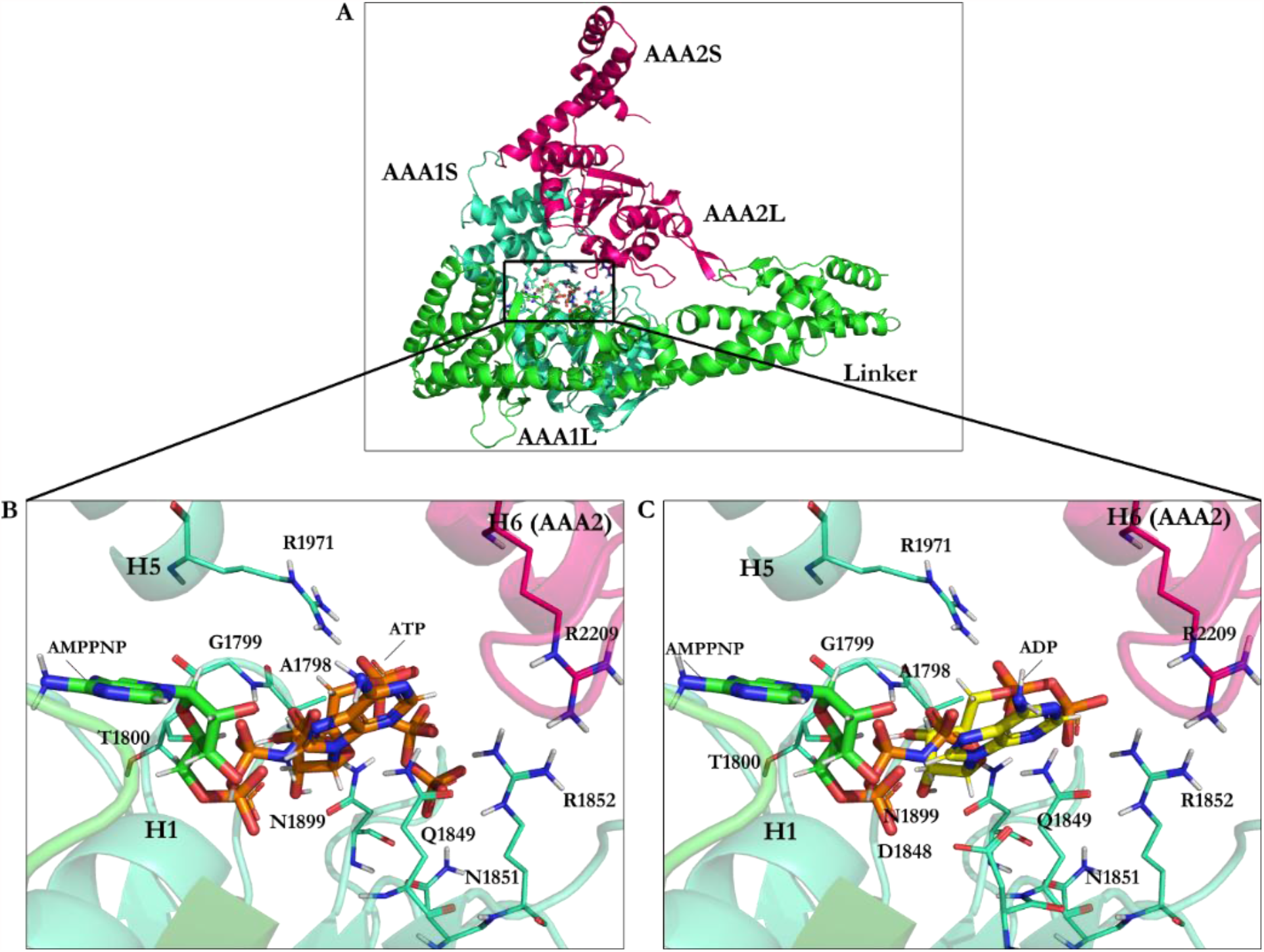
**A)** The linker, the AAA1, and the AAA2 subunits of dynein and illustration of their inter-subunit binding site. Binding interactions of docking solutions of **B)** ATP (orange) and **C)** ADP (yellow) compared to AMPPNP (green) at the AAA1 binding site.

The ADP’s binding energy is -31.89 kJ/mol, which is the highest among the nucleotides (ATP with -42.33 kJ/mol and AMPPNP with -40.18 kJ/mol), thus presenting the lowest affinity towards the AAA1 binding site. **(Figure 7 C & Table 5)**

#### 3.2.1. Ciliobrevin A and D

The calculated conformation of ciliobrevin A (binding energy -26.23 kJ/mol) had a stronger affinity than ciliobrevin D (binding energy -23.92 kJ/mol). Ciliobrevin A (IC_50_ of 52.0 µM [26]) has a lower potency than the D analog (IC_50_ of 15.0 µM [26]). Ciliobrevin A and D both display weaker binding affinity than ATP (−42.33 kJ/mol), AMPPNP (−40.18 kJ/mol), and ADP (−31.89 kJ/mol). **(Figure 4 & Tables 5-6)**

The O21 atom of ciliobrevin A is involved in a ∼ 2.1 Å hydrogen bond (H-bond). In contrast, the O21 of ciliobrevin D forms a ∼ 1.8 Å H-bond with Lys1802 (the W-A motif, Gly1796-Thr1803); the positively charged ammonium fragment of Lys1802 usually contributes to the stabilization of the negatively charged ATP γ-phosphate [37]. The O22 of ciliobrevin A and D engage in H-bond with the side chain of Asn1899 S-I motif of the β6 (Ala1893-Asn1899) from ∼ 1.3 Å -1.4 Å distance. The S-I is involved in placing a water molecule near the γ-phosphate of ATP and the negative charge of Glu1849 of the W-B motif, thereby facilitates a nucleophilic attack for hydrolyzation [37]. The O22 in ciliobrevin A and D also the formation of an H-bond (∼ 1.8 Å and ∼ 1.9 Å, respectively) with Gln1849 (in E1849Q mutant). In the wild-type dynein, Glu1849 is responsible for activating a water molecule placed by the S-I (Asn1899 in yeast dynein 1) to trigger the network mechanism of ATP hydrolysis [37]. Therefore, the E1849Q mutation in the yeast-motor-AMPPNP crystal structure represents a conformation incapable of ATP hydrolysis [20]. In the *in silico* conformational search that the mutant of dynein was studied, Gln1849 showed interactions with ciliobrevin A and D by H-bonds formation. The N9 atom of the ligands interacted with the hydroxyl group of Thr1803 of the W-A motif (Gly1796-Thr1803) through a 2.8 Å H-bond in ciliobrevin A, and a 2.9 Å H-bond in ciliobrevin D, while Thr1803 usually participates in the stabilization of the ATP γ-phosphate [37]. Thus, by interacting with Thr1803, ciliobrevin A and D could block the activity of the subsites, which otherwise would be involved in the hydrolytic reaction on the ATP. **(Figure 4 & Figure 8)**

**Figure 8:**
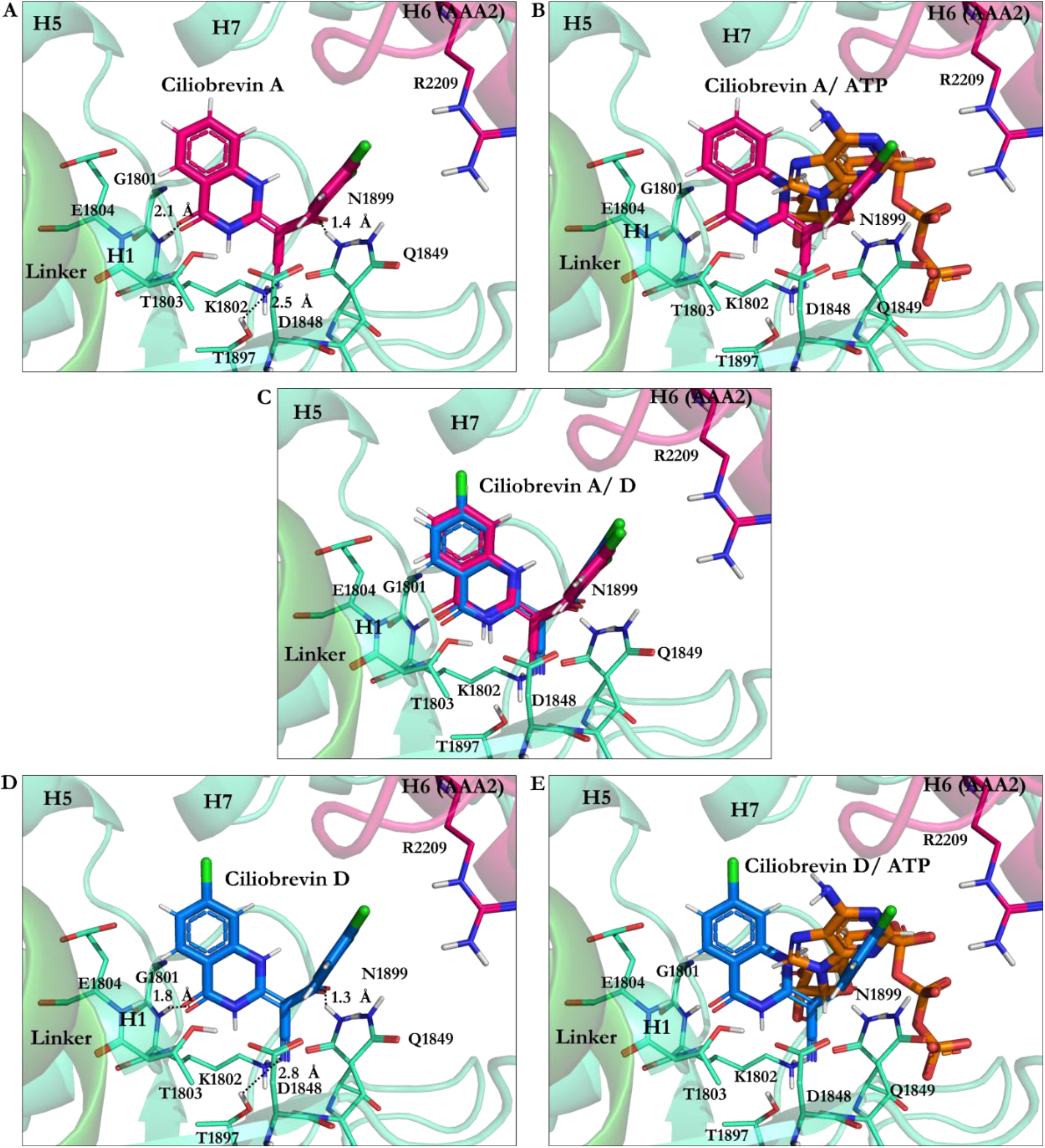
Ciliobrevin A and D conformation at the AAA1 binding site of motor domain of dynein 1. Ciliobrevin A, **B)** Ciliobrevin A superimposed on ATP, **C)** Ciliobrevin A and ciliobrevin D superimposed, **D)** Ciliobrevin D, **E)** Ciliobrevin D superimposed on ATP.

The cyanide (CN) moiety of ciliobrevin A has an H-bond (∼ 2.5 Å) with Thr1897 of β6 (Ala1893-Asn1899). The CN is involved with the hydroxyl (OH) moiety of Thr1897 in ciliobrevin D (from ∼ 2.8 Å distance). Noteworthy that Thr1897 does not belong to the ATP motifs, nor does it show any interactions in the *in silico* docking solutions of the nucleotides. However, the CN moiety seems to act as an auxiliary anchor to promote placements and orientations of the significant substructures of ciliobrevin A and D in the proximity of the critical ATP motifs, namely the W-A and the S-I. Aliphatic chains of the W-A (by Gly1799, Lys1802), the S-I (by Asn1899), and the W-B motif (by Asp1848 and Gln1849) are involved in van der Waals (VdW) interactions with the hydrophobic fragments of ciliobrevin A, and of ciliobrevin D. That suggests how ciliobrevin A and D’s effects as ATP antagonists, on the ATP motifs and Thr1897 could disturb the activity of the motor domain by blocking the catalytic residues action. **(Figure 8 &Table 3)**

#### 3.2.2. The Analogs Binding Profile

Analog 30 showed the best affinity among analog 29 and 28 (the lowest energy -27.87 kJ/mol, vs. respective -27.37 kJ/mol and-27.27 kJ/mol). The O21 in analogs 28, 29, and 30 is involved in an H-bond with the polar H of the amide bond moiety of Gly1801 in the ATP motif, the W-A (Gly1796-Thr1803). Furthermore, their N9 atom forms H-bond with the OH moiety of Thr1803 of the W-A (Gly1796-Thr1803). In these analogs, the CN moiety plays a similar role in ciliobrevin A and D. It is also involved in an H-bond formation with the OH of Thr1897 in the β6. The side chains of Asn1899 in the S-I and Gln1849 of the W-B motif also create H-bond with the O22 of the analogs. The hydrocarbon chains of Gly1799 and Lys 1802 in the W-A motif, and Asp1848 as well as Gln1849 of the W-B motif, hydrophobically interact with the C8 of quinazolinone ring B and the acrylonitrile moiety. Thus, the analogs 28-30 engage with ATP motifs and the β6, through which they could hinder the motor domain’s natural function. **(Figures 5, Figure 9 & Table S1)**

**Figure 9:**
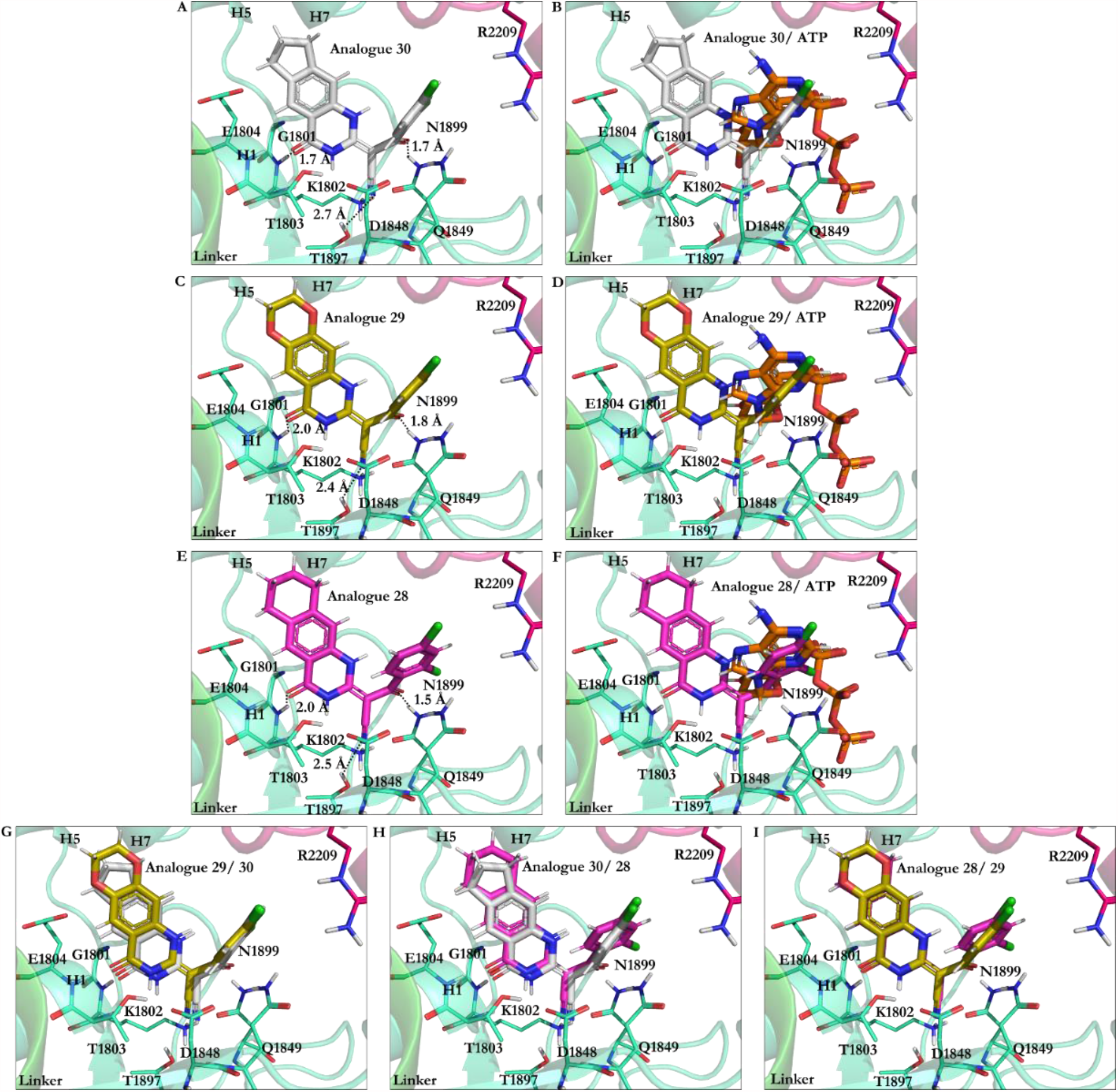
Analogs of ciliobrevin at the AAA1 binding site of dynein 1. **A)** Analog 30, **B)** Analog 30 superimposed on ATP, **C)** Analog 29, **D)** Analog 29 superimposed on ATP, **E)** Analog 28, **F)** Analog 28 superimposed on ATP, Superimposition of **G)** Analogs 29 and 30, **H)** Analogs 30 and 28, and **I)** Analogs 28 and 29.

Chemical modifications resulting in analog 45 showed its improved binding affinity versus analogs 28-30, 38, and 42, ciliobrevin A and D. However, the similarity in their binding profiles showed they could comparably compete with ATP for binding to the functional motifs in the AAA1 nucleotide - binding site. The analogs O21 atom forms H-bond with the polar H of the Gly1801 the W-A motif, and their N9 atom with the OH of Thr1803 in the same ATP motif. There is also an H-bond between the O22 and Asn1899 of the S-I, and Gln1849 of the W-B. Similar to the 28-30, the CN moiety of analog 42 forms H-bond with Thr1897 of the β6. Thr1897, which does not belong to the ATP motifs, interacts with ciliobrevin A and D, analogs 45, 30, 42, 29, 28 and 38, as well. The OH of Thr1897 is involved with Gln1849 (in the W-B motif), known for its connection with water molecules to promote ATP hydrolysis. In analog 38, the CN was replaced with a methoxy (-OMe) moiety. The O34 of the methoxy group interacts with Lys1974 via a 2.2 Å H-bond. The benzene ring was replaced with pyridine in analog 45, whose N31 atom forms a 2.1 Å H-bond with Lys1974 in the H7 helix (Leu1970-Pro1982). The Lys1974 positive charge could potentially form a dipole-induced moment with the pyridine ring of analog 45, although the positively charged amino acid is not perpendicular to the ring. **(Table S1-2, Table 3 & Figure S6)**

The analogs of ciliobrevin (i.e., ciliobrevin A and D, 28, 29, 30, 38, 42, and 45) affect the ATP hydrolysis process also through binding to Asn1899 (in the S-I), which normally forms H-bond with and position water molecules for the nucleophilic substitution [37]. A conserved Asn residue (e.g., Asn64 in PspF, a member of the AAA+ proteins [38]) is involved in an H-bond formation with the conserved Glu from the W-B motif (Glu108 in PspF of AAA+ proteins [38]) found at the AAA1 binding site of several dyneins [37]. Through the interactions of an ATP competitive inhibitor with the Glu or the Asn, the Asn (Asn64 in PspF [38]) could not contribute to the ATP hydrolysis, as the glutamate residue of the W-B motif (Glu108 in PspF [38]) is unavailable to activate a water molecule through deprotonation [37]. The process is referred to as a “glutamate switch” and thought to be an endogenous mechanism that regulates ATP hydrolysis in dynein to evade non-productive powerstroke [37]. H-bond formations of Asn1899 with ciliobrevin A, D and its analogs could disrupt the “switch”-mechanism and therefore interfere with the regulation of the dynein powerstroke progression. The glutamate switch involving Glu108 in PspF [38] has not yet been detected in cytoplasmic dynein 1. The “glutamate switch”-involving Asn residue is replaced with a cysteine (Cys1822 in *S. cerevisiae*) in the dynein 1 isoform [37]. However, an intramolecular H-bond of 2.1 Å between the side chains of Gln1849 (E1849Q) and Arg1852 was visualized in the optimized (i.e., energy minimized) structure of yeast-motor-AMPPNP dynein obtained through an *in silico* conformational search. In contrast, the crystal structure shows a relatively long distance (3.9 Å) between the residues. Thus, the energetically stabilized conformation demonstrates Arg1852 and Gln1849 in the positions and orientations capable of strong H-bond formation, where an arginine in place of the asparagine could interact with the glutamate to execute the “switch” mechanism in dynein 1. **(Figure S7)**

#### 3.2.3. Geometrical Isomerization Effect on Ciliobrevin Binding to the AAA1

Ciliobrevin A and D exist in two geometric isomers of *E* or *Z* at the C8-C11 double bond [10]. The potency of the ciliobrevin was thought to be affected by isomerization, where there is only a fraction of the isomer abrogating dynein [4]. The benzoylacrylonitrile group of the molecule favors the *E* isomer since the one-dimensional NMR spectrum and the result of a 2D Nuclear Overhauser Effect Spectroscopy (NOESY) of ciliobrevin D, has shown an intramolecular H-bond between the hydrogen atom on the N7 and the O22 that stabilizes ciliobrevin D in solution [26]. The N7 is directly attached to the C8, and the O22 relates to the C11 via the double bond to the C13, which is covalently attached to the C11. **(Figure 5 & Figure S8)**

*Ciliobrevin A:* The effect of geometrical isomerization of ciliobrevin A was investigated, where its *Z* isomer showed binding with 6.82 kJ/mol higher energy than of the *E* isomer (−26.23 kJ/mol). The O22 atom of the *Z* isomer engaged in a 1.9 Å H-bond with the side chain of the S-I, through Asn1899 in β6 (Ala1893-Asn1899) and a 2.7 Å H-bond with β3 (Ala1843-Asp1848) of the W-B via Gln1849. The *Z* isomer did not interact with Thr1897 of the β6 (Ala1893-Asn1899), unlike the *E* isomer of ciliobrevin A. Whereas, its CN moiety forms an H-bond with the S-II ATP motif through Arg1971 of the H7 helix (Leu1970-Pro1982). The N9 atom of the *Z* isomer involved in a 2.2 Å H-bond with the carboxylate moiety of Asp1848 from β3 (Ala1843-Asp1848), and its O21 atom also formed a 2.2 Å H-bond with the amino group of Asn1821 in the β2 strand (Val1818-Asn1821). Ring A of the *Z* isomer oriented to form a π-π stacking with the guanidine moiety of Arg1852, a key element in the “glutamate switch” in dynein 1, as observed in the conformation obtained in this *in silico* conformational search. The binding energies of the *E* and *Z* isomers of ciliobrevin A indicate that the *E* is favorable over the *Z* isomer, as the former displayed a significantly higher binding affinity towards the AAA1 binding site. **(Figure 10 A-B & Table S3)**

**Figure 10:**
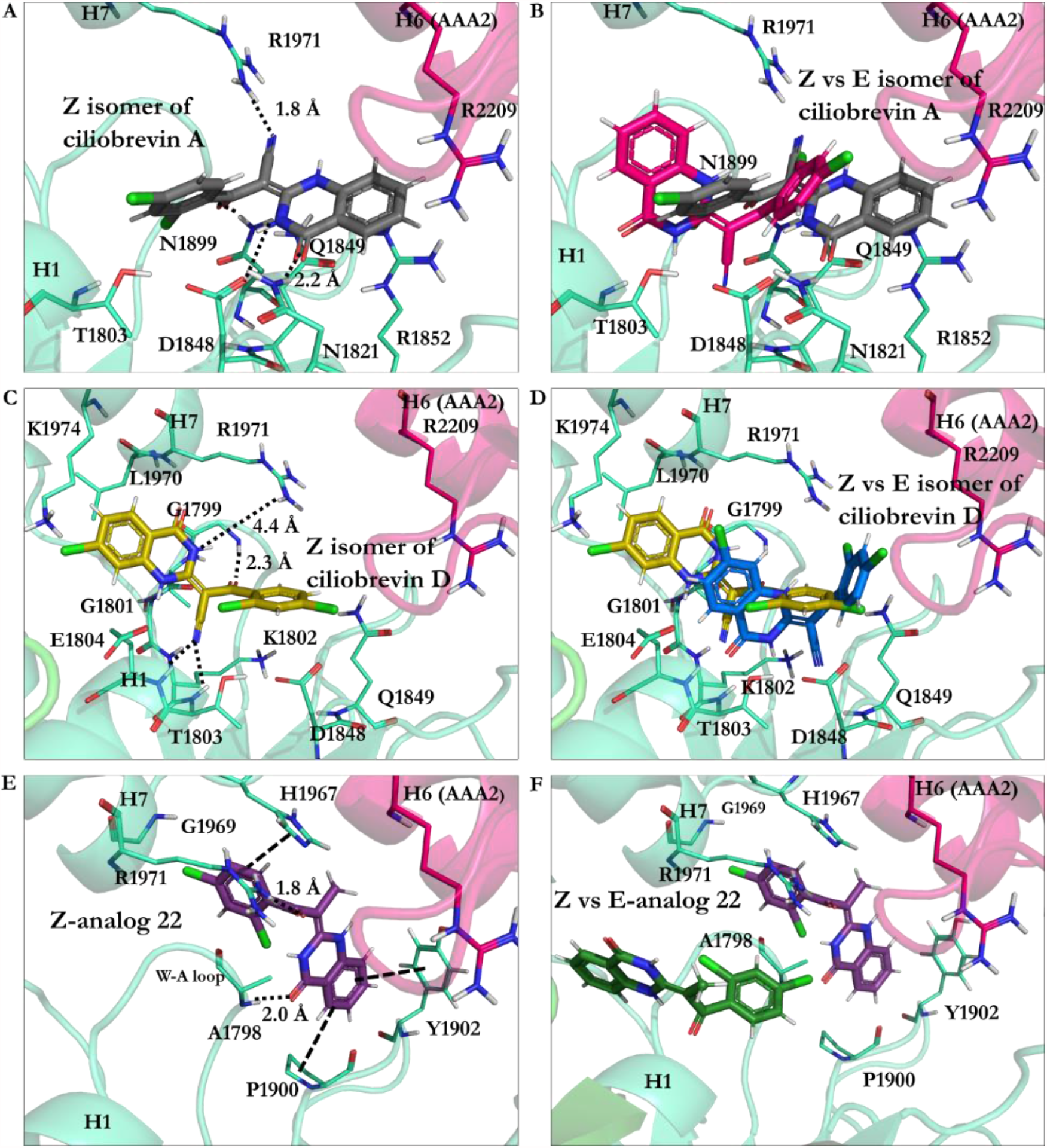
**A)** *Z* isomer of ciliobrevin A at the AAA1 binding site of cytoplasmic dynein 1, **B)** Superimposition of *Z* and *E* isomers of ciliobrevin A, **C)** *E* isomer of ciliobrevin D at the AAA1 binding site of cytoplasmic dynein 1, **D)** *E* isomer of ciliobrevin D superimposed on its *Z* isomer, **E)** *Z*-analog 22 and the residues at the AAA1 binding site of AAA1, **F)** *Z*-analog 22 superimposed on its *E* isomer at the AAA1 binding site.

*Ciliobrevin D:* The binding energy of its *Z* isomer, similar to ciliobrevin A, was also higher than its *E* (- 19.78 kJ/mol vs. -23.92 kJ/mol, respectively). The *Z* isomer utilized its O22 atom to engage in an H-bond with the polar hydrogen of the Gly1799 amide moiety in the W-A motif (Gly1796-Thr1803). The *Z’*s CN group was available to form H-bonds with Lys1802 (1.9 Å) of the W-A motif, Thr1803 (2.5 Å), and Glu1804 (2.3 Å). The N9 of the *Z* isomer has a weak H-bond with Arg 1971 in the H7 helix (Leu1970-Pro1982). Noteworthy that the *Z* isomer of ciliobrevin D does not interact with the S-I motif via Asn1899. **(Figure 10 C-D & Table S3)**

The effect of the CN elimination from ciliobrevin A and its replacement with a methyl group in analog 22 [7] was examined through the study of its geometric isomers. The *E* isomer became weaker than *E*-ciliobrevin A and D; however, slightly stronger than its *Z* isomer (analog 22 *E* isomer, -17.49 kJ/mol, versus -16.83 kJ/mol). The *E* is bound to the AAA1 site via H-bond with Lys1802. It also formed an H-bond via its N9 atom with the carboxylate group of Glu1804. The *Z* isomer of analog 22 displayed a ∼ 180 ° rotation in the binding site compared to its *E* isomer and ciliobrevin A and D. Its unique orientation caused H-bonds with Ala1798 and Arg1971 via its O22 atom. In addition, His1967 interacted with its ring D of the *Z* isomer through a T-shaped π-π stacking. Its ring A also showed a similar conformation against Tyr1902, while the Pro1900 orientation facilitated a proline-benzene VdW interaction via the ligand’s ring D. **(Figure 10 F & Table S3)**

#### 3.2.4. Dynapyrazoles A and B

The O21 of dynapyrazole A and B forms H-bonds (∼1.9 Å) with the W-A peptide backbones (Gly1796-Thr1803) via Lys1802 and H1 helix (H1, Glu1804-Gly1810). The H atom on the N9 in dynapyrazole A and B are involved in a 1.6 Å H-bond with the W-A via Thr1803, whose OH moiety typically interacts with an Mg2+ resulting in stabilizing the charges on the ATP γ-phosphate [37]. Thus, the ligands, which have a slight binding difference (∼ 0.32 kJ/mol), could similarly hinder the interaction between the cation and the Thr. In addition, the CN moiety of dynapyrazole A and B forms H-bonds with the β2 strand (Val1818-Asn1821) through Asn1821 (2.3 Å), which is ∼4.0 Å far from Asp1848, a member of the W-B motif in the β3 (Ala1843-Asp1848). They could indirectly impact dynein’s motility affecting the W-B’s Asp1848, a segment that usually hosts ATP to undergo hydrolysis [4, 37]. **(Figure 11 & Table S4)**

**Figure 11:**
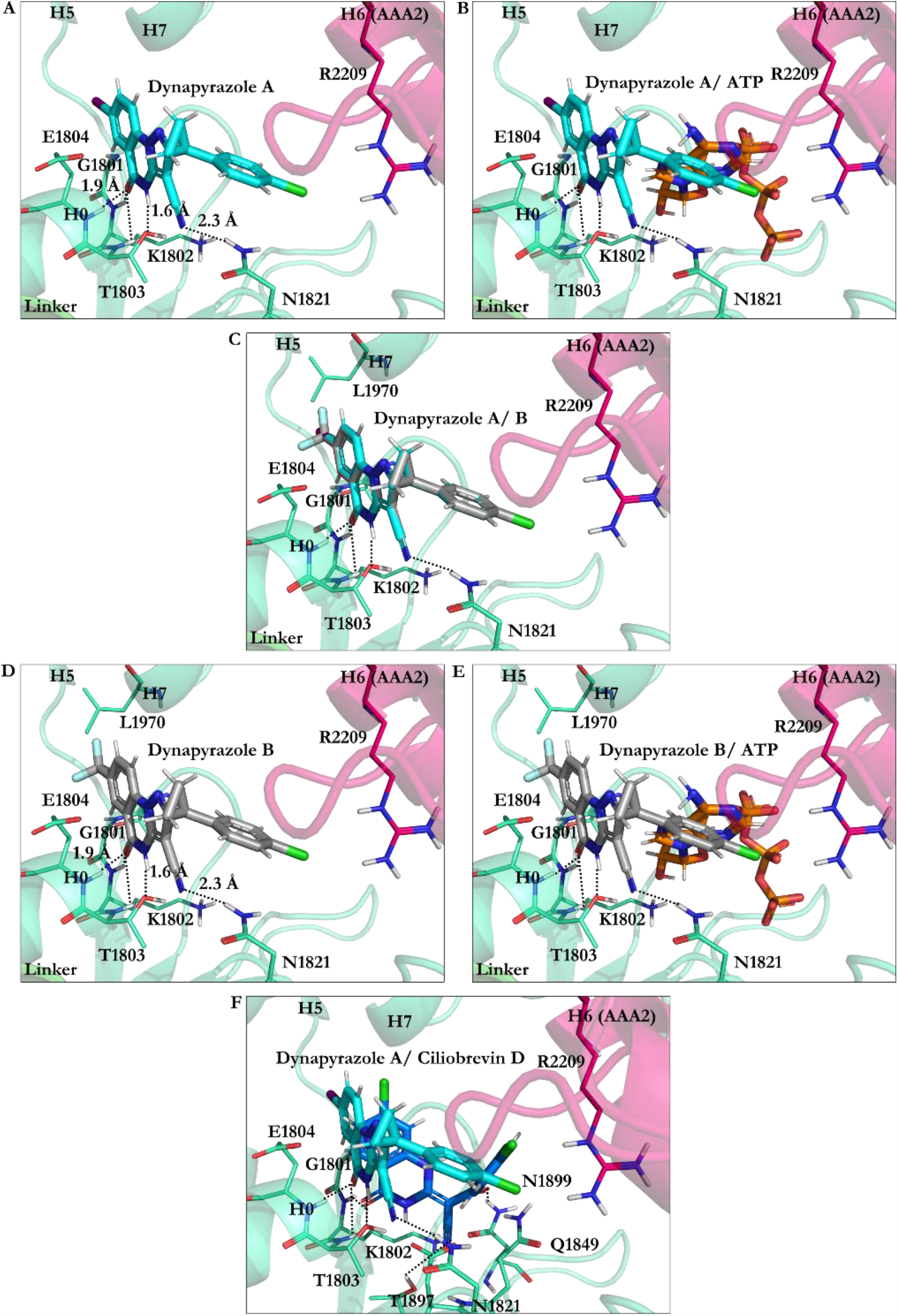
Dynapyrazole in the nucleotide-binding site of the AAA1 **A)** dynapyrazole A, **B)** Superimposition of dynapyrazole A and ATP, **C)** Superimposition of dynapyrazole A and B, **D)** Binding modes of dynapyrazole B, **E)** Superimposition of dynapyrazole B and ATP, **F)** Superimposition of dynapyrazole A and ciliobrevin D.

#### 3.2.5. Impact of Elimination of Carbon Double Bond on Dynapyrazole and Analogs Affinity

Ciliobrevin’s derivatization led to the synthesis of dynapyrazole by eliminating the C8-C11 double bond and inserting the ring C in dynapyrazole and its analogs [26]. The process also consisted of replacing the O22 atom in ciliobrevin with the N11 in dynapyrazole to improve its potency. That resulted in the IC_50_ plummet from 15 µM (ciliobrevin D) to 2.3 µM (dynapyrazole A) [26], whereas the binding strength of the former improved for ∼ - 5 kJ/mol. **(Figure 4 & Table 4, Table S3-S4)** The double bond in ciliobrevin A and D positionally allows the *E* isomer to form H-bonds both with Thr1897 of the β6 (Ala1893-Asn1899) via the nitrogen of its CN moiety and with Asn1899 through its O22 atom. The energy contribution of this event could be the cause of the difference in the total binding strength, considering that the N11 of the replaced-ring C in dynapyrazole, has no interaction with the AAA1 binding site, in contrast to the eliminated O22 in ciliobrevin. However, the CN nitrogen atom of dynapyrazole A and B forms an H-bond with Asn1821. **(Figure 11 F)** Among the analogs 37, 43, and 47 of ciliobrevin possessing the double bond, analog 47 shows the lowest IC_50_ (130.0 µM [26]) and the strongest binding (−26.12 kJ/mol), whereas analog 37 with the highest IC_50_ (280.0 µM [26]) has just 2.01 kJ/mol higher binding energy than analog 47. The analog 47 is suggested as the most suitable candidate for further *in vitro* and *in vivo* experimental evaluations for its effect on dynein motility and its selectivity profile. **(Figure 12)**

**Figure 12:**
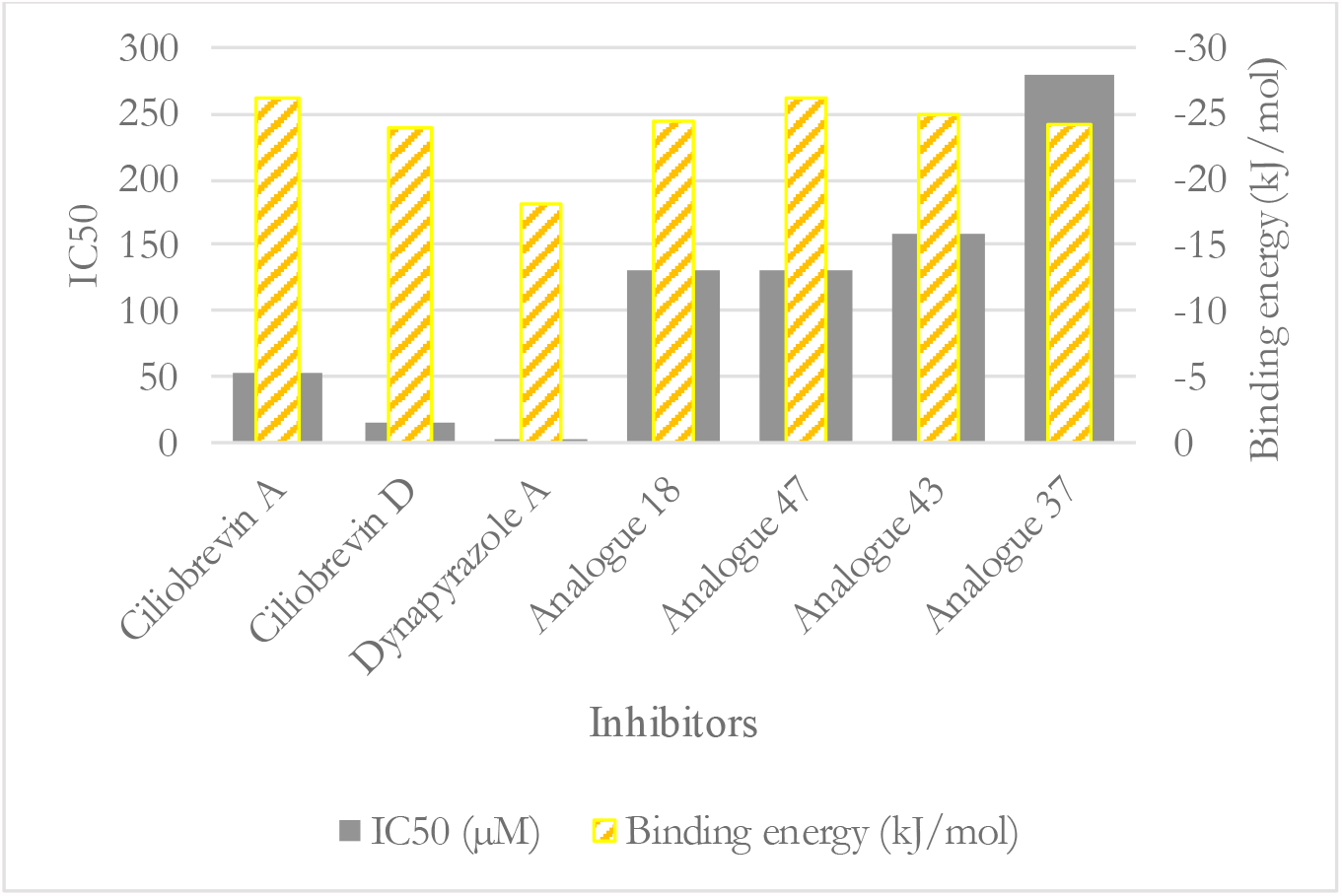
The ligands binding energies versus their IC_50_.

#### 3.2.6. Protonation Effect on Ciliobrevin A and D Binding

Considering the pKa values of the chemical moieties of the inhibitors as stated earlier, the N9 and N7 might be weak candidates for protonation at the tissues with alkaline pH. **(Figure 4, Figure S9 & Table 6)**

**Table 6:** Amino acids affected by ciliobrevin A and D in their protonated and deprotonated states.

| Compound | Binding energy<br>(kJ/mol) | Residues interacting with the compound |
| --- | --- | --- |
| **Ciliobrevin A N9**<br>**protonated** | -28.22 | Ala1798, Lys1802, Thr1803, Glu1804, Asp1848,<br>Gln1849, Thr1897, Asn1899, Arg1971 |
| **Ciliobrevin A** | -26.23 | Gly1801, Lys1802, Thr1803, Glu1804, Asp1848,<br>Gln1849, Thr1897, Asn1899 |
| **Ciliobrevin D N9**<br>**protonated** | -26.15 | Lys1802, Thr1803, Glu1804, Asp1848, Gln1849,<br>Thr1897, Asn1899 |
| **Ciliobrevin D** | -23.92 | Gly1801, Lys1802, Thr1803, Glu1804, Asp1848,<br>Gln1849, Thr1897, Asn1899 |
| **Ciliobrevin A N7**<br>**protonated** | -23.84 | Gly1799, Thr1800, Gly1801, Lys1802, Thr1803,<br>Glu1804, Leu1970, Arg1971 |
| **Ciliobrevin D N7**<br>**protonated** | -23.55 | Gly1801, Lys1802, Thr1803, Glu1804, Asp1848,<br>Gln1849, Thr1897, Asn1899 |

##### The protonation of ciliobrevin A at the N9 position

caused a ∼ -2.0 kJ/mol improvement in binding strength and suggesting that ciliobrevin A might be protonated at the N9 depending on the environmental pH when interferes with the motor function. However, the possibility seems low concerning the juxtaposed carbonyl moiety at the C10. Unlike the neutral (unprotonated) ciliobrevin A, its protonated form interacts with Ala1798 of the W-A motif (Gly1796-Thr1803) through its benzylic ring D. The protonated N9 atom is 2.6 Å far from the carboxylate moiety of Asp1848 in the β3 (Ala1843-Asp1848), which could also electrostatically affect the positively charged N9 atom and contribute to the strengthening of ciliobrevin A binding affinity.

##### The protonated N9 of ciliobrevin D

projects a similar binding profile to that of the A analog with an enhanced binding compared to its neutral form (−23.92 kJ/mol vs. -26.15 kJ/mol). Therefore, ciliobrevin D, protonated under a proper pH, has a superior inhibitory effect on dynein 1 *in vitro*.

##### The protonated ciliobrevin A at the N7

is a weaker binder (∼ 2.39 kJ/mol) compared to its neutral structure (with -26.23 kJ/mol), suggesting the ligand is less likely to be protonated at the N7 position in solution, as the nitrogen atom’s lone pair-electrons tend to participate in the delocalized electron cloud of the aromatic ring A.

##### The protonation of the N7 atom of ciliobrevin D

had a minor effect on binding (0.37 kJ/mol), since protonated and unprotonated D analog similarly treat the AAA1 nucleotide site through Gly1801, Lys1802, Thr1803, Glu1804, Asp1848, Gln1849, Thr1897, Asn1899.

#### 3.2.7. Effect of Protonation on Binding Mode of Dynapyrazole A and B

##### Protonation of dynapyrazole A at N9

resulted in a slight binding improvement (∼ - 0.64 kJ/mol). This analog is the only one in the library of 63 ligands to bind to the linker domain of dynein. An ionic interaction is formed between the protonated N9 and the carboxylate group of Glu1586 in the T-turn 6 (Val1586-Glu1588 of the linker), while its O21 has H-bond with the amide moiety of Pro1766 in the N-loop (Pro1766-Leu1774). The aliphatic side chains of Lys1696 and Glu1699 in the H13 of the linker domain (Asp1692-Asn1717) and Glu1767 in the N-loop (Pro1766-Leu1774) are hydrophobically affected by the hydrocarbon fragment of the ligand consisting of the C8, the C12, and the C13 atoms. Positively charged guanidium moiety of Arg1978 in the H7 (Leu1970-Pro1982) binds the monochloride benzylic ring D through polar interactions. The observations elucidate the improvement of the total binding strength of the N9-protonated dynapyrazole A.

##### Protonation of dynapyrazole A at N7

also benefits from protonation (∼ - 8.72 kJ/mol). The considerable improvement indicates the IC_50_ *in vitro* better correlates with the ligand’s protonated at the N7 position. It electrostatically interacts with the negative charge of Glu1804 carboxylate in the H1 helix (Glu1804-Gly1810), while its N9 creates an H-bond to the OH moiety of Thr1803 in the W-A (Gly1796-Thr1803). The data showed that protonation at the N7 site is beneficial to the ligand. **(Figure 4, Figure 13 & Table S4)**

**Figure 13:**
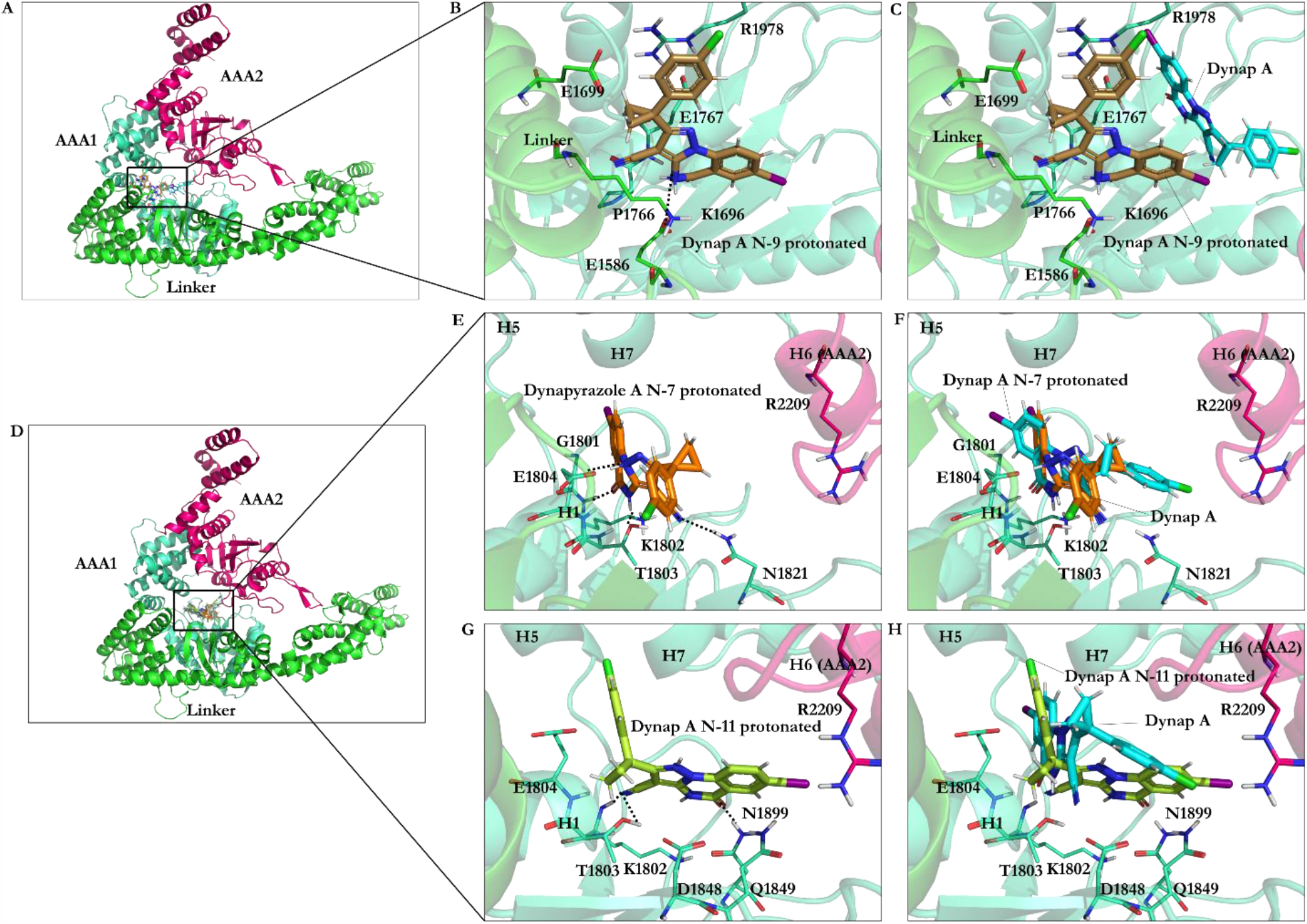
Binding the protonated dynapyrazole A at the AAA1 binding site of dynein 1. **A)** Overview of the AAA1, AAA2 units, and linker domains, **B)** Dynapyrazole A protonated at the N9 atom interacting with the linker residues, **C)** Dynapyrazole A protonated at the N9 atom superimposed on dynapyrazole A, **D)** Overview of the AAA1, AAA2, and linker subdomains (as the panel A), **E)** Dynapyrazole A protonated at the N7 atom, **F)** Dynapyrazole A protonated at the N7 atom superimposed on dynapyrazole A, **G)** Dynapyrazole A protonated at the N11 atom, and **H)** Dynapyrazole A protonated at the N11 atom superimposed on dynapyrazole A.

##### Protonation of N11 in dynapyrazole A

caused slight weakness of the binding (0.27 kJ/mol) compared to its neutral form due to a minor difference of the interaction-network set up by the N11-protonated ligand. The protonated N11 atom shows no ionic interactions. Dynapyrazole A in this configuration is the only analog, among the protonated and neutral dynapyrazole A and B, to interact with the β6 (via Asn1899) and the β3 strand (via Asp1848 and Gln1849), resembling the binding mode of ciliobrevin and its analogs.

##### Protonation of dynapyrazole B at N11

also had an insignificant effect on its binding (∼ 0.62 kJ/mol), similar to its protonated A analog. It has the lowest predicted affinity towards the AAA1 subunit in the ligands library and involves Val1819 through VdW forces via its ring D. In summary, protonation at the N11 is disadvantageous to dynapyrazole A and B and weakens their binding affinities to the AAA1 site. **(Figure 4, Figure S10 E-F & Table S4)**

## 4. Conclusion

The presented work provides structural data according to a SAR study to explain how ciliobrevin A, D, and dynapyrazole A, B, their protonated structures as well as the forty-six analogs could inhibit ATP binding and its hydrolysis in the nucleotide-binding site of the AAA1 subunit of the motor domain in cytoplasmic dynein 1. The lowest binding energy of ATP among the 63 ligands of the library suggested its superior binding affinity over all the competitive inhibitors. However, ciliobrevin A, D, and most of the analogs bind to the functionally key sub-sites, including the Sensor I and II, N-loop, and the W-A and B, also known as the ATP motifs; thus, optimizing the concentration of the competitive inhibitors *in vitro* could result in blocking the AAA1 nucleotide site in the absence of the ATP or its lower concentration. In particular, analog 47 is suggested as the most suitable candidate for further *in vitro* and *in vivo* experimental evaluations due to its strong binding affinity and low IC_50_. The ligands’ structural mechanism of interference with the ATP binding and hydrolysis was shown to vary depending on their critical functional fragments. The presence of carbonyl oxygen on the ring B of the ligands, for instance, in ciliobrevin D, resulted in its O21 atom involving an H-bond with Lys1802 amine moiety in the W-A motif. The positively charged ammonium group of the Lys usually acts as an anchor by applying electrostatic forces on the negatively charged γ-phosphate, thereby contributing to the catalytic network for ATP hydrolysis. At the same time, the O22 of the ligand forms an H-bond with Asn1899 of the S-I motif in the β6 strand. The S-I is involved in placing a water molecule near the γ-phosphate of ATP and the negative charge of Glu1849 of the W-B motif and enables a water molecule for a nucleophilic attack, required for the ATP hydrolyzation. The O22 also facilitated the ligand’s hydrogen bond formation with the Gln1849 in the Glu1849Gln protein mutant.

Eliminating the C8-C11 double from the ciliobrevin, removing the O22, and replacing it with the N11 by insertion of the ring C in dynapyrazole, resulted in the alteration of the chemical structure, which lowered the IC_50_ in dynapyrazole. However, the N11 of the ring did not mimic the O22 effect and diminished dynapyrazole binding strength, despite being at a relatively similar position in the ligand structure. Protonation at the N11 atom did not enhance its contribution to the binding energy, as shown in a separate attempt. However, dynapyrazole A benefits from the N7 and N9 atoms protonation according to the improvement gained in their binding energy. The N9 protonated dynapyrazole A is the only analog in the ligand library, which binds to the linker domain of dynein. The ligand conformational pose and the consequent binding to the linker domain were facilitated by the electrostatic interaction between the protonated N9 and the carboxylate group of Glu1586 in the T-turn 6 of the linker and an H-bond with the amide moiety of Pro1766 in the N-loop. The aliphatic side chains of the H13 helix in the linker domain, as well as the N-loop, interacted with the ligand hydrophobic sites, namely, the C8, the C12, and the C13 atoms. The observation explained the improvement of the total binding strength of the N9-protonated dynapyrazole A against its unprotonated form.

There are two geometrical isomers *E* and *Z* of ciliobrevin, according to its C8-C11 double bond. The *E* isomer enabled H-bond formation of the ligand with the β6 via its Thr1897 through the nitrogen of the ligand CN moiety and with Asn1899 via its O22 atom. The *Z* isomer of the analog D interacted with the W-A motif, showing no substantial effect on the S-I motif. In contrast, the *Z* isomer of ciliobrevin A interacted with the S-I motif through Asn1899 of the β6, the β3 strand, and with the W-B via Gln1849. Unlike its *E* isomer, the *Z* of ciliobrevin A showed no effect on Thr1897 of the β6. However, its CN moiety caused an H-bond with the S-II motif. The ring A, benzene moiety of the *Z* isomer, had a polar interaction with Arg1852 in a position suitable for a π-π stacking with the guanidine moiety of the arginine. It also appears to contribute to the “glutamate switch” mechanism in dynein 1. The binding energies of the geometric isomers of ciliobrevin A and D indicated that the *E* had a significantly higher affinity than the *Z* towards the AAA1 binding site. Assessing the effect of geometrical isomerization on analog 22 resulted in the conformation of its isomers in two opposite orientations in the binding site. This was likely due to the replacement of the CN moiety with a methyl group in that particular analog, causing a drastic change in the isomers binding modes.

The benzene ring replaced with a pyridine moiety in analog 45 has a polar interaction with Lys1974. The analog 45, similar to other pyridine possessing analogs of ciliobrevin (i.e., 28, 29, 30, 38, 42, and 45) could affect the ATP hydrolysis via binding to Asn1899, a conserved residue of the S-I motif that facilitates placing water molecules for nucleophilic substitution in the ATP hydrolysis.

The glutamate switch (involving Glu108 in PspF [38]) has not yet been detected in cytoplasmic dynein 1; however, an intramolecular H-bond of 2.1 Å between the side chains of Gln1849 (E1849Q) and Arg1852 was detected in the most energetically favorable conformation of the yeast-motor-domain through the *in silico* conformational search. It exhibited a conformation with Arg1852 and Gln1849 in orientations capable of a close and strong H-bond formation, suggesting that arginine could also interact with the glutamate in regulating the “switch” mechanism in dynein 1.

New analogs of dynapyrazole, recently introduced by Santarossa *et al*.,[36] have shown to be potent in inhibiting basal ATPase activity of dynein while binding to AAA3 and AAA4. The binding assessment of analogs 3-48 will be critical in evaluating their inhibitory mode of action at the AAA3 binding site, which is not conserved in axonemal dynein and cytoplasmic dynein 2. This makes the AAA3 subunit a suitable target in the future direction of this study. Utilizing the presented information contributes to setting up the experiments to focus on the most potential analogs for their selectivity to dynein 1 versus its 2-isoform.

## Supplementary Information

**Figure S1:** Amino acids sequence alignment of the AAA1 and AAA2 subunits of dynein corresponding to *S. cerevisiae* (1758-2273) and *D. discoideum* (1936-2531) with the ClustalW program. The W-A motif is highlighted in blue. The grey highlights represent sequence similarity. A singular dot (.) indicates that residues belong to a different group of amino acids, while a colon (:) Residues belong to the same group of amino acids and the asterix (*) indicates sequence identity.

**Figure S2: A)** Structural alignment of the AAA1 and AAA2 of yeast-AMPPNP (4W8F) and yeast-apo (4AKG), **B)** A close-up of the AAA1 binding site. Helices (H0, H1, H5, and H7) of the AAA1 subdomain, the Walker-A (W-A) loops with the AMPPNP in the ball and stick representation in green located at the AAA1 nucleotide-binding site and the H6 of the AAA2 subdomains of the yeast-AMPPNP (4W8F) and yeast-apo (4AKG). Color code: yeast-AMPPNP (AAA1 in green cyan and AAA2 in sharp pink) and yeast-apo (AAA1 in orange and AAA2 in yellow).

**Figure S3: A)** Structural alignment of the AAA1 and AAA2 subunit of the Dictyostelium-ADP (3VKG) and yeast-AMPPNP (4W8F), **B)** A close-up of the AAA1 binding site. Helices (H0, H1, H5, and H7) of the AAA1 subdomain, the Walker-A (W-A) loops with the AMPPNP in the ball and stick representation in green and ADP in the ball and stick representation in yellow located at the AAA1 nucleotide-binding site and the H4 of the AAA2 subdomain of the Dictyostelium-ADP (3VKG) and the H6 of the AAA2 subdomain of the yeast-apo (4AKG), **C)** Structural alignment of ligands at the AAA1 nucleotide-binding site of the Dictyostelium-ADP (3VKG) and yeast-AMPPNP (4W8F) with AMPPNP in the ball and stick representation in green and ADP in the ball and stick representation in yellow with a cross (x) on its atoms to distinguish them from the atoms of AMPPNP. Color code: Dictyostelium-ADP (AAA1 in green and AAA2 in blue) and yeast-AMPPNP (AAA1 in green cyan and AAA2 in sharp pink).

**Figure S4: A)** AMPPNP interacting and residues of the AAA1 nucleotide-binding site. Distances between catalytic residues of the AAA1 nucleotide-binding site of dynein and the β-phosphate of **B)** AMPPNP and **C)** ADP.

**Figure S5:** Annotation of the secondary structure of AAA1 and AAA2 domains of cytoplasmic dynein according to the 4W8F crystal structure retrieved from the PDB.

**Figure S6:** Binding modes of ciliobrevin analogs at the AAA1 binding site of dynein 1. A) Analog 45. Analog 45 superimposed on ATP, C) Analog 42, D) Analog 42 superimposed on ATP, E) Analog 38, F) Analog 38 superimposed on ATP. Superimposition of G) Analogs 42 and 45, H) Analogs 38 and 45, I) Analogs 38 and 42.

**Figure S7:** Intramolecular interactions between the catalytic residues at the AAA1 binding site and Arg1852 involving a potential glutamate switch. The docking solution of AMPPNP is represented in green and stick. Thr1803 and Asp1848 accommodate Mg2+ during ATP hydrolysis. Sensor I (N1899), W-B (E1849Q), R-finger (R2209) play catalytic roles during ATP hydrolysis.

**Figure S8:** *E* and *Z* configurations of ciliobrevin D. Red arrows point to the double bond between the C8 and C11 atoms.

**Figure S9:** Binding modes of the protonated forms of ciliobrevin at the AAA1 binding site of dynein 1. **A)** Ciliobrevin A protonated at the N9 atom, **B)** Ciliobrevin A protonated at the N9 atom superimposed on ciliobrevin A, **C)** Ciliobrevin A protonated at the N7 atom, **D)** Ciliobrevin A protonated at the N7 atom superimposed on ciliobrevin A, **E)** Ciliobrevin D protonated at the N9 atom, **F)** Ciliobrevin D protonated at the N9 atom superimposed on ciliobrevin D, **G)** Ciliobrevin D protonated at the N7 atom, **H)** Ciliobrevin D protonated at the N7 atom superimposed on ciliobrevin D.

**Figure S10:** Main binding interactions of the protonated forms of dynapyrazole B at the AAA1 binding site of dynein 1. **A)** Dynapyrazole B protonated at the N7 atom, **B)** Dynapyrazole A protonated at the N7 atom superimposed on dynapyrazole B, **C)** Dynapyrazole B protonated at the N9 atom, **D)** Dynapyrazole B protonated at the N9 atom superimposed on dynapyrazole A, **E)** Dynapyrazole B protonated at the N11 atom, **F)** Dynapyrazole B protonated at the N11 atom superimposed on dynapyrazole B.

**Table S1:** Networking interactions of analogs 28, 29, and 30 with AAA1 of dynein 1.

**Table S2:** Interacting amino acids with analogs 38, 42, and 45.

**Table S3:** The binding energy of *E* and *Z* isomers of ciliobrevin A and ciliobrevin D.

**Table S4:** Interacting amino acids with dynapyrazole A and B in the protonated and deprotonated states.

## Acknowledgments

The authors acknowledge funding support from the Natural Sciences and Engineering Research Council of Canada (NSERC), Discovery Grant (No. 212654), awarded to L.A. The authors thank Compute Canada for the technical supports provided through the ACNET team.

## Associated Content

The authors will release the atomic coordinates and experimental data upon article publication.

## Corresponding Author

Laleh Alisaraie*

## Author Contributions

The manuscript was written by the contributions of the authors. The authors have approved the final version of the manuscript.

